# Protective *IFIH1* variant reduces immune-mediated islet stress and dysfunction in a type 1 diabetes genetic background

**DOI:** 10.64898/2025.12.30.697107

**Authors:** Daniel A. Veronese-Paniagua, Cameron Banks, Kameron Bradley, Noyonika Mukherjee, Sarah E. Gale, Kate E. Hinshaw, Amy M. Meacham, Opeoluwa F. Iwaloye, Hubert M. Tse, Clayton E. Mathews, Jeffrey R. Millman

## Abstract

Genome-wide association studies (GWAS) have linked dozens of genetic loci to type 1 diabetes (T1D). The *IFIH1* gene, which encodes the double-stranded RNA sensor MDA5, is one such locus. The E627* single nucleotide polymorphism (SNP) in *IFIH1* is associated with protection against T1D, while the A946T variant is linked to increased risk. While the E627* variant has been shown to result in a truncated protein and dampen type I interferon (IFN) signaling, its specific role in human pancreatic islet health and function remains unclear. We hypothesized that MDA5^627^* would protect islet cells from stress-induced dysfunction, identity loss, and cell death. Using CRISPR-Cas9 technology, we introduced the E627* and A946T variants into human pluripotent stem cells (hPSCs) derived from a T1D patient. We differentiated these hPSCs into stem cell-derived islets (SC-islets) and treated them with IFNα, poly(I:C), and coxsackievirus B3, an enterovirus implicated in T1D pathogenesis. Using single-cell RNA sequencing and an array of functional assays, we investigated the variant impact on both whole SC-islets and their individual cell populations. Our analysis revealed that SC-islets, and their β, α, and δ cell subpopulations, harboring the MDA5^627^* variant exhibit an attenuated immune response to the various stressors compared to MDA5^946T^ cells. We also report unique, cell-type-specific transcriptional responses that vary across variants. Notably, MDA5^627^* SC-islets showed reduced apoptosis rates and viral genome expression, as well as attenuated negative effects on mitochondrial function and insulin secretion in response to stress. Overall, our findings demonstrate that a clinically relevant MDA5 variant confers protection by dampening stress-mediated transcriptional responses, reducing cell dysfunction, and preventing apoptosis. These insights provide a mechanistic framework for understanding T1D pathogenesis and offer new avenues for developing preventative therapies.

## Introduction

Type 1 diabetes (T1D) is a CD8+ T cell-mediated autoimmune disease characterized by the progressive destruction of insulin-secreting β cells within the pancreatic islets of Langerhans. This destruction results in little to no insulin secretion and subsequent high blood sugar levels in patients. Affecting 27-54 million people worldwide with a rising incidence rate, T1D presents a significant global health challenge.^1^ Current treatments, such as daily insulin injections and pancreatic islet transplantation, have limitations. Insulin injections fail to precisely mimic the complex regulatory mechanisms of islets, leading to dangerous fluctuations in blood sugar and long-term complications.^2–5^ While islet transplantation offers a functional cure, it requires multiple organ donors, strict immunosuppression, and is susceptible to eventual graft failure.^6–8^ Therefore, widening our understanding of T1D initiation and progression is critical to developing preventative therapies.

Genetic predisposition is a major driving factor in T1D pathology. Studies attribute roughly 50% of the genetic risk to Major Histocompatibility Complex (MHC)/human leukocyte antigen (HLA) and *INS* genes.^9–12^ Furthermore, genome-wide association studies (GWAS) have identified over 70 non-HLA loci associated with T1D.^11,13–17^ However, genetic background is not the sole factor involved in disease onset. The rising incidence of T1D in immigrant populations moving to high-incidence countries and a 35% discordance rate in monozygotic twins highlight the critical role of environmental factors.^18,19^ The seasonal pattern of T1D onset also suggests a potential trigger from external exposures, leading researchers to investigate factors such as viral infections.^20–23^

Viral infections, especially by enteroviruses like coxsackievirus B (CVB), have long been implicated as a potential environmental trigger for T1D.^9,24^ Enteroviral RNA has been detected in the stool of T1D-susceptible individuals and in the pancreatic islets of recent-onset patients.^25,26^ Moreover, clinical trial data have shown that antiviral therapy can help preserve residual insulin production in T1D patients.^27^ This link between viral infection and T1D is strongly supported by the association of T1D with genes involved in antiviral immune response, such as Interferon Induced with Helicase Domain 1 (*IFIH1*).^10,13,17,28^ The *IFIH1* gene encodes Melanoma Differentiation-Association Gene 5, an RNA helicase that detects cytoplasmic double-stranded RNA (dsRNA) to trigger a type I interferon (IFN)-mediated antiviral immune response.^29–31^

GWAS studies have linked various single nucleotide polymorphisms (SNPs) within *IFIH1* to T1D risk.^32^ Studies utilizing peripheral blood mononuclear cells (PBMCs) from T1D patients have shown that the *IFIH1* A946T variant (*rs1990760*) is a risk allele that strengthens the type I IFN response to stress.^33,34^ In mice, it also increased T1D incidence.^33,35^ Conversely, the E627* variant (*rs35744605*), which introduces an early termination codon, is associated with a dampened immune response in PBMCs and protection against T1D.^36–38^ While studies using PBMCs and mouse models have provided valuable insights, the role of these *IFIH1* variants in human pancreatic islet cell health and function has been difficult to determine. The scarcity of islets from T1D patients, combined with their inherent dysfunction, presents a significant research gap.

Human pluripotent stem cells (hPSCs) have proven to be a valuable tool for modeling diabetes.^39,40^ Many research groups have developed protocols using growth factors and compounds to mimic human pancreatic development, generating stem cell-derived pancreatic islets (SC-islets).^41–50^ These SC-islets are composed of over 80% pancreatic endocrine cells, can secrete insulin, and have been shown to cure T1D when transplanted into diabetic mice.^41,42,45^ Additionally, SC-islets address several limitations associated with animal models, such as differences in islet architecture and genetics.^51,52^ They also overcome the issues presented by cadaveric islets, namely donor-to-donor genetic variability and a limited supply. Since SC-islets can be generated as an unlimited supply of genetically identical cells, they increase experimental robustness and expand the capabilities for genetic studies. As a result, SC-islets have enabled groups to model islet development using hPSCs from T1D donors.^53,54^ They have also allowed for the investigation of how genetic candidates and diabetes-associated genes influence β cell development and overall islet health, highlighting their potential for studying diabetes pathology.^39,40,53–67^

In this study, we hypothesized that SC-islets harboring a protective *IFIH1* variant have a dampened immune response that protects β cells from stress-induced dysfunction, identity loss, and death. To test this hypothesis, we utilized CRISPR-Cas9 edited hPSCs reprogrammed from a T1D patient to generate SC-islets. We investigated the role of the protective E627* *IFIH1* variant in islet stress response by treating SC-islets with IFNα, the synthetic double-stranded RNA polyinosinic-polycytidylic acid (poly(I:C)), and CVB3. We utilized single-cell RNA sequencing to delineate whole islet and cell type-specific transcriptional responses to the different treatments. We found that the MDA5^627^* variant leads to a dampened immune response in all three SC-endocrine cell types, while also differentially impacting their transcriptional responses. We validated our findings with functional assays, measuring mitochondrial function, insulin secretion, and cell death. Our data support a connection between *IFIH1* SNPs, stress, and β cell health, highlighting the utility of this dataset in delineating the impacts of *IFIH1* genetic variants on islet health in a human context. These findings also further validate SC-islets as a robust *in vitro* model for studying clinically relevant genetic variants in the context of diabetogenic viral infections.

## Results

### Generation and characterization of human SC-islets harboring different T1D-associated IFIH1 variants

To investigate the effects of a protective *IFIH1* variant in a T1D genetic background, we used the CH1-064 hPSC line, which the New York Stem Cell Foundation reprogrammed from a 59-year-old Caucasian female with a T1D diagnosis at age 51. We used CRISPR-Cas9 to edit the *IFIH1* gene at either *rs1990760* (A946T) or *rs35744605* (E627*) in the parental line to generate MDA5^946T^ and MDA5^627^* edited hPSCs, respectively (Fig. 1a, Extended Data Fig. 1a-g, Supplementary Data 1). Compared to the parental line, there was no significant change in *IFIH1* gene expression in MDA5^946T^ hPSCs (Fig. 1b, Supplementary Data 2). However, in MDA5^627^* hPSCs, *IFIH1* expression was less than half of that measured in the parental line.

**Figure 1.**
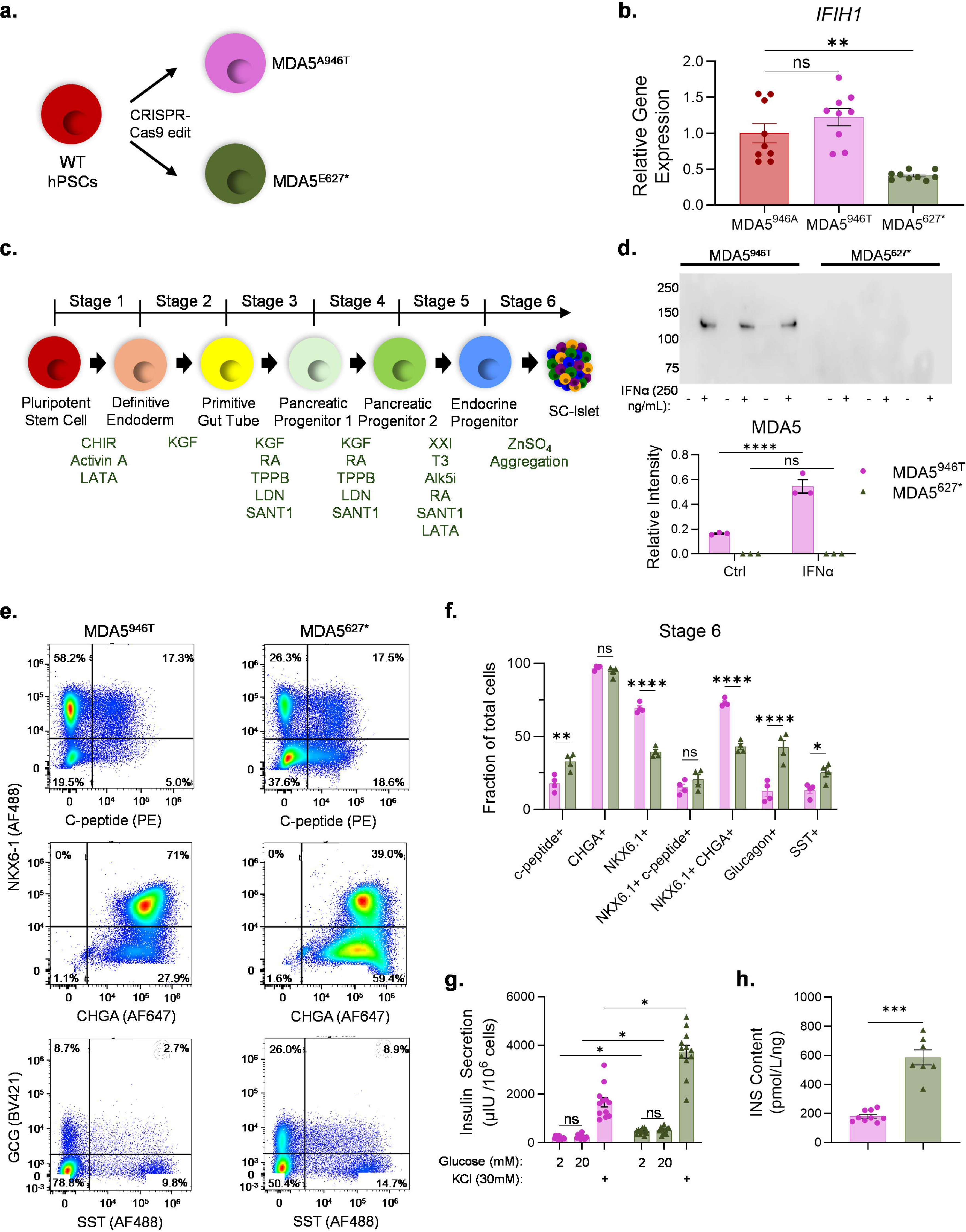
Generation and characterization of *IFIH1* mutant SC-islets. **(A)**, Schematic depicting the generation of *IFIH1* genetic variants. **(B)**, rt-qPCR of *IFIH1* expression in parental and CRISPR-Cas9 edited hPSC lines (n = 3). ** = Brown-Forsythe and Welch ANOVA tests followed by Dunnett’s multiple comparisons test. **(C)**, Schematic of SC-islet differentiation protocol adapted from Hogrebe et al. (2025). **(D)**, Representative and quantification of western blot of MDA5 expression in SC-islets following treatment with 250ng/mL IFNα (n = 3). ** = ** = Brown-Forsythe and Welch ANOVA tests followed by Dunnett’s multiple comparisons test. **(E)**, Representative flow cytometry dot plots of dispersed SC-islets generated from each MDA5 variant for NKX6-1, C-peptide, CHGA, GCG, and SST. **(F)**, Quantified fraction of cells expressing or co-expressing islet markers for SC-islets derived from each MDA5 variant (n = 4). ** = Two-way ANOVA followed by Sidak’s multiple comparisons test. **(G)**, Bar graphs measuring static glucose-stimulated insulin secretion of SC-islets (n = 4). ** = Multiple unpair t tests and two-stage linear step-up procedure of Benjamini, Krieger, and Yekutieli. **(H)**, Bar graphs measuring intracellular insulin content in SC-islets from each variant (n = 3). ** = Welch’s unpaired t test.

To study *IFIH1* variants in a pancreatic context, we adapted a previously reported SC-islet differentiation protocol with minor modifications to the Latrunculin A concentrations in stage 1 (Fig. 1c, Supplementary Data 2).^41,42,68^ We attempted to differentiate the parental line into SC-islets but were unable to obtain a high yield of SC-endocrine cells, as most of the population consisted of off-target cell types (data not shown). In contrast, the differentiation of both MDA5^946T^ and MDA5^627^* hPSCs was successful. To determine if the MDA5 protein was properly expressed in SC-islets, we treated the SC-islets with a high concentration of IFNα to induce high expression of MDA5 and performed western blot using an antibody targeting the residues around Arg470 within the Helicase 1 domain of the protein. Our analysis revealed increased MDA5 expression following treatment in MDA5^946T^ SC-islets but no detectable protein expression in both untreated and treated MDA5^627^* SC-islets (Fig. 1d).

In the SC-islets from both cell lines, over 90% of the population was composed of endocrine-like cells, marked by the co-expression of NKX6-1 and CHGA (Fig. 1e-f). The generation of SC-β cells, as demonstrated by co-expression of NKX6-1 and C-peptide, was lower in MDA5^946T^-derived cells (15.06%) when compared to MDA5^627^* SC-islets (20.55%), though this difference was not statistically significant (Fig. 1e-f). However, MDA5^946T^-derived SC-islets did have significantly lower fraction yields of SC-α (12.29% vs 42.35%) and SC-δ (13.19% vs 25.45%) cells, as marked by GCG and SST expression, respectively (Fig. 1e-f). Additionally, we observed significantly higher proportions of an endocrine, intestinal-like off-target population, termed SC-enterochromaffin (EC) cells, in MDA5^946T^ (69.86%) when compared to MDA5^627^* SC-islets (36.85%) (Extended Data Fig. 2a-b).

Next, we assessed the secretory function of the SC-islets from each variant. A static glucose-stimulated insulin secretion (sGSIS) assay confirmed that the SC-islets from both *IFIH1* variants could release insulin when challenged with glucose (Fig. 1g). However, there was no statistical difference in insulin release within each variant when switched from 2mM to 20mM glucose. The addition of the membrane depolarizer, KCl, resulted in increased insulin secretion. We noted that the insulin secretion response to both glucose and KCl was stronger in MDA5^627^* SC-islets (Fig. 1g). We also observed higher intracellular insulin content in MDA5^627^* SC-islets compared to MDA5^946T^ SC-islets (Fig. 1h).

### Single-cell transcriptomics reveals a divergent stress response in the MDA5^627^* SC-islets

To determine if the MDA5^627^* variant protects pancreatic cells from stress, we differentiated hPSCs into SC-islets and then treated them for 24 and 48 hours (h) with water (control), IFNα, the synthetic double-stranded RNA polyinosinic-polycytidylic acid (poly(I:C)), or CVB3-Woodruff (Fig. 2a). CVB3 has been shown to have a strong affinity for pancreatic tissue and to accelerate T1D onset in non-obese diabetic (NOD) mouse models.^67,69,70^ Considering that 3D cultured cells require higher concentrations than their 2D counterpart, we infected whole SC-islets with a multiplicity of infection (MOI) of 20, which is consistent with previous studies on primary human islets.^67,71–76^ After treatment, we fixed SC-islets and processed them for single-cell RNA sequencing (scRNA-seq). Following quality check and filtering, we analyzed 113,322 cells across all conditions and time points (Extended Data Fig. 3a). Cell marker gene expression confirmed the presence of α, β, and δ cells in both MDA5 variants (Fig. 2b-c, Extended Data Fig. 3b-c). We identified five EC cell subpopulations, two of which expressed multiple stress-associated genes, including *DDB2*, *TRIAP1*, *ATF3*, and *NFKBIA*. We also observed clusters containing endocrine progenitors, identified by positive expression of *INS*, *GCG*, and *SST* and a lack of *PDX1*, *NKX6-1*, and *HES1* gene expression (Fig. 2c, Extended Data Fig. 3c). Proliferating cells formed a separate cluster marked by high expression of cell cycle and cell division-associated genes, such as *MKI67* and *CDC6*. Sequencing results were consistent across batches, demonstrating the robustness of SC-islets as a disease model (Extended Data Fig. 3d).

**Figure 2.**
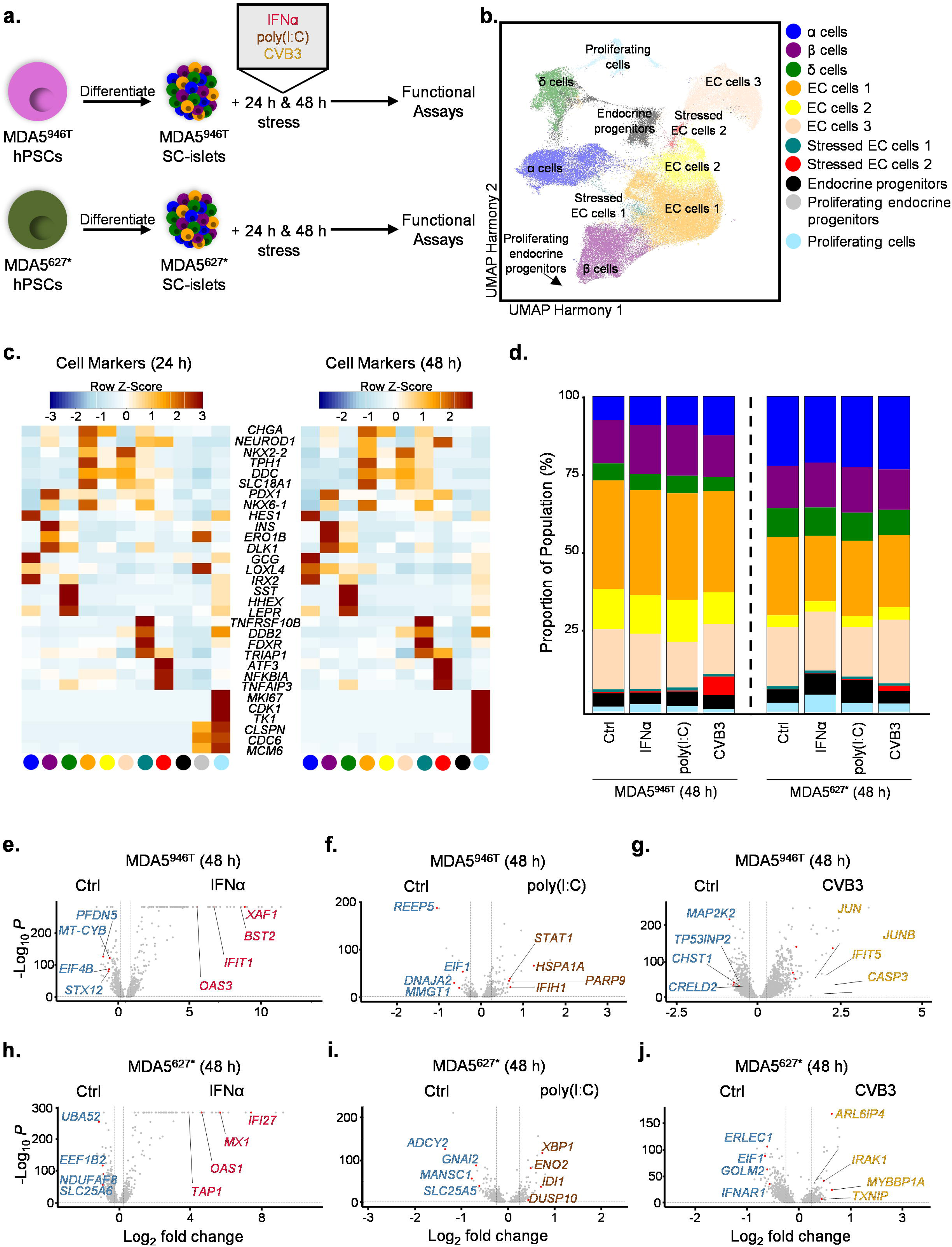
scRNA-seq of SC-islets reveals transcriptional differences between MDA5 variants following stress treatments. **(A)**, Workflow schematic for treatment processing of SC-islets. **(B)**, Harmony UMAP of cell populations identified following scRNA-seq data integration. **(C)**, Heatmaps depicting the z-score of genetic marker expression per cluster for each time point. **(D)**, Stacked bar plots depicting the proportion of cells found in SC-islets derived from each MDA5 variant and treated with various stressors for 48 h. **(E-J)**, Volcano plots showing transcriptional differences when comparing **(E)** MDA5^946T^ Ctrl vs IFNα 48 h (total variables = 1920), **(F)** MDA5^946T^ Ctrl vs poly(I:C) 48 h (total variables = 769), **(G)** MDA5^946T^ Ctrl vs CVB3 48 h (total variables = 3326), **(H)** MDA5^627*^ Ctrl vs IFNα 48 h (total variables = 1700), **(I)** MDA5^627*^ Ctrl vs poly(I:C) 48 h (total variables = 1083), and **(J)** MDA5^627*^ Ctrl vs CVB3 48 h (total variables = 1815).

All clusters were identified in both variants except for proliferating endocrine progenitors, which were only present in CVB3-infected MDA5^946T^ SC-islets processed 24 h after treatment (Fig. 2c, Extended Data Fig. 4a-b). Although stress treatment did not impact the proportion of cell types within SC-islets from either *IFIH1* variant, the cell proportions were not similar across them at both time points (Fig. 2d, Extended Data Fig. 4b, Supplementary Data 3). For instance, at 48 h, MDA5^946T^ control SC-islets exhibited lower proportions of SC-α cells (7.48% vs 21.96%) and higher proportions of all identified EC cells together (67.41% vs 48.32%). SC-β cells proportions were similar (13.83% vs 13.36%) across both variants in this control group. These trends were consistent across time points and treatments (Fig. 2d, Extended Data. Fig. 4b).

We performed a pseudo-bulk analysis to identify differentially expressed genes (DEGs) by conducting pairwise comparisons between control and treatment conditions at each time point. At 24 h and 48h, IFNα-treated MDA5^946T^ SC-islets differentially expressed 2,066 and 1,920 genes, respectively, with the upregulation of multiple type I IFN-associated genes, including *OAS2*, *IFITM3*, *IFIT1*, and *BST2* (Fig. 2e, Extended Data Fig. 4c, Supplementary Data 4). In contrast, control expressed genes associated with protein synthesis, transcription, and protein folding (*EEF2*, *EIF4B*, *PFDN5*).

At 24 h, poly(I:C) treatment of MDA5^946T^ SC-islets induced differential expression of 382 genes, such as *BEX2* (regulates cell cycle and apoptosis), *POM121* (a component of the nuclear pore complex), and *REPS1* (a signaling adaptor protein), while 48 h saw 769 DEGs, including immune-associated genes (*STAT1*, *PARP9)* (Fig. 2f, Extended Data Fig. 4d, Supplementary Data 4). When compared to poly(I:C) at both time points, MDA5^946T^ control cells expressed genes linked to protein synthesis, transcription, and protein folding, including *EEF2, EIF4B*, *PFDN5*, *DNAJA2*, and *XBP1*.

CVB3 infection of MDA5^946T^ SC-islets induced 1,197 and 3,326 DEGs at 24 h and 48 h, respectively (Fig. 2g, Extended Data Fig. 4e, Supplementary Data 4). At 24 h, CVB3 infection upregulated *MOB1B* (a regulator of Hippo signaling), *ODC1* (involved in polyamine biosynthesis and cell growth), and the tumor suppressor *ARRDC3*, while control SC-islets had higher *PRPF8* (a spliceosome component), *ANKS6* (associated with cilia), and *CBX6* (a chromatin regulator) expression. *JUN*, *JUNB*, *IFIT5*, and *CASP3*, all associated with cellular immune response, were upregulated following CVB3 infection, while *MAP2K2*, *CRELD2*, and *CHST1* were observed in control cells.

We observed 1,880 and 1,700 DEGs in MDA5^627^* SC-islets treated with IFNα for 24 and 48 h, respectively, as well as increased expression of multiple type I IFN-associated genes, such as *ISG15*, *IFIT3*, *IFI27*, *OAS1*, and *MX1* at both time points (Fig. 2h, Extended Data Fig. 4f, Supplementary Data 4). In contrast, control cells showed higher expression of *SRP9* (a protein transport regulator) and *RAB4A* (a protein trafficking regulator) at 24 h, and *NDUFAF8* (a mitochondrial component) and *SLC25A6* (a mitochondrial carrier) at 48 h.

In MDA5^627^* cells, poly(I:C) treatment resulted in 940 DEGs at 24 h and 1,083 DEGs after 48 h (Fig. 2i, Extended Data Fig. 4g, Supplementary Data 4). Poly(I:C)-treated SC-islets increased *RTRAF* (associated with transcription and translation), *SERF2* (regulates protein destabilization), and *BUB3* (involved in cell division) at 24 h, while we observed increased expression of stress-associated *XBP1*, *ENO2*, and *DUSP10* in the 48 h condition. Compared to poly(I:C) treatment, control cells expressed *SRP9*, *ISCU* (a component of the iron-sulfur cluster scaffold), and *BEX2* after 24 h, while the 48 h control cells upregulated *ADCY2* (a key regulator in cyclic adenosine monophosphate production), *MANSC1* (associated with the Golgi Apparatus [Golgi]), and *SLC25A5* (an ADP-ATP antiporter).

We observed 872 and 1,815 DEGs in CVB3-infected MDA5^627^* SC-islets after 24 and 48 h, respectively (Fig. 2j, Extended Data Fig. 4h, Supplementary Data 4). Upregulated genes at 24 h following infection included *RTRAF*, *FUS* (an RNA binding protein), and *ITPR3* (involved in calcium signaling), while control cells expressed *COA3* (involved in mitochondrial complex assembly), *FUNDC1* (associated with autophagy), and *PCGF3* (a chromatin remodeler). *ARL6IP4* (associated with mRNA splicing) and stress-associated genes *IRAK1* and *TXNIP* were upregulated in CVB3-infected cells after 48 h. Control cells expressed *IFNAR1* (type I IFN receptor), *EIF1* (associated with transcription), and *GOLM2* (associated with the Golgi).

We performed a pairwise comparison between the two *IFIH1* variants at basal conditions to better understand their transcriptional differences. Our analysis revealed that MDA5^946T^ and MDA5^627^* differentially expressed a total of 3,983 genes (Extended Data Fig. 4i, Supplementary Data 4). SC-islets harboring the truncated MDA5^627^* exhibited higher expression of islet identity genes, including *GCG* and *INS*, as well as *JPH3* (associated with the endoplasmic reticulum) and *DPP6* (binds potassium channels). In contrast, MDA5^946T^ overexpressed stress-associated genes, such as *NALP2*, *ISG15*, and *HSPA2*.

### Differential gene expression is governed by cell type and IFIH1 variant

To leverage the advantages of scRNA-seq, we performed pairwise comparisons between treatment and control groups for each cell type in our dataset. The analysis revealed that the transcriptional response to the stressors is cell-type specific and differs across variants (Fig. 3a-b, Supplementary Data 5-6).

**Figure 3.**
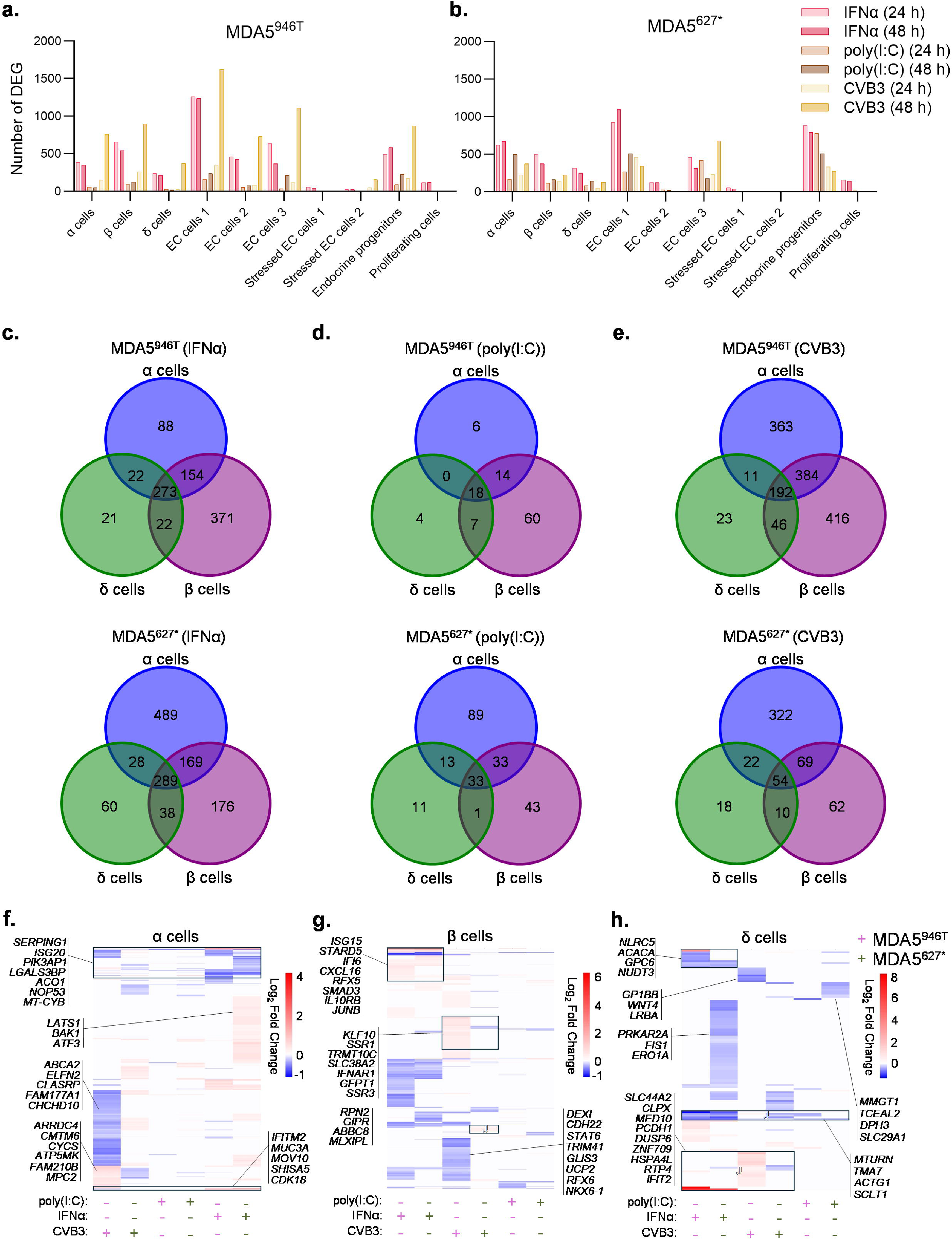
scRNA-seq of SC-islets reveals major cell type-specific transcriptional differences between SC-endocrine cells and MDA5 variants. **(A-B)**, Bar graphs showing the number of differentially expressed genes (DEGs) for each cell type found in **(A)** MDA5^946T^ and **(B)** MDA5^627*^ SC-islets. **(C-E)**, Venn diagrams comparing DEGs from both time points for SC-α, -β, and -δ cells from each MDA5 variant treated with either **(C)** IFNα, **(D)** poly(I:C), or **(E)** CVB3. **(F-H)**, Heatmaps showing log_2_ fold change of DEGs unique to **(F)** SC-α, **(G)** SC-β, and **(H)** SC-δ cells from each MDA5 variant following stress treatments.

MDA5^627^*-derived SC-α and SC-δ cells showed a larger number of DEGs across all time points and treatments, with the exception of the 48 h CVB3 infection, when compared to their MDA5^946T^ counterparts (Fig. 3a-b). In contrast, MDA5^946T^ SC-β cells induce more DEGs when treated with IFNα and CVB3 across both time points but not following poly(I:C) treatment. We also determined that MDA5^627^*-derived pancreatic endocrine cell types were most transcriptionally responsive to IFNα treatment, with two-fold or higher number of DEGs compared to the next most impactful stressor, CVB3. In contrast, MDA5^946T^-derived endocrine cells were most sensitive to IFNα at 24 h and CVB3 at 48 h (Fig. 3a-b). Given that SC-EC are derived from an intestinal lineage and that CVB3 infection commences in the gastrointestinal tract in humans,^9^ we also compared the transcriptional response across variants. Overall, SC-EC cells generated from MDA5^946T^ hPSCs exhibited a larger transcriptional response to stress treatment. Additionally, the majority of DEGs corresponding to both stressed EC cell clusters arose in MDA5^946T^ cells, with no DEGs in MDA5^627^* cells following CVB3 infection and only five in total after poly(I:C) treatment (Fig. 3a-b). MDA5^627^* endocrine progenitors and proliferating cells induced stronger transcriptional responses across all conditions and time points, except for 48 h of CVB3 infection. Considering our focus on the effects of the MDA5^627^* variant on pancreatic endocrine cells, we excluded all other cell types from downstream analyses.

### Stress treatment induces both conserved and unique transcriptional signatures in SC-endocrine cells

We next assessed the unique transcriptional responses of each endocrine cell type to the various stressors and investigated their similarities across MDA5 variants. To do this, we combined the data from both time points and performed pairwise comparisons of treatment and control conditions for each cell type.

IFNα-treated SC-α, -β, and -δ cells shared 273 and 289 DEGs in MDA5^946T^ and MDA5^627^* SC-islets, respectively (Fig. 3c, Supplementary Data 7). When considering the DEGs unique to each cell type, we observed that MDA5^946T^ SC-β differentially regulated more than two-fold the number of DEGs compared to MDA5^627^* SC-β cells and 4.22-17.67-fold more than SC-α and -δ cells also harboring the MDA5^946T^ variant. However, the number of unique DEGs for MDA5^627*^ SC-α cells was more than four-fold when compared to their counterpart in MDA5^946T^ and 2.78-8.15-fold higher than MDA5 ^627*^ SC-β and -δ cells (Fig. 3c). MDA5^627*^-derived SC-δ cells had nearly three-fold more DEGs than MDA5^946T^ SC-δ cells. The number of DEGs shared between pairs of cells was roughly similar in both variants.

Poly(I:C) treatment resulted in 18 shared DEGs within the three MDA5^946T^ cell types and 33 DEGs within the MDA5^627*^ cells (Fig. 3d, Supplementary Data 7). Similar to the trend with IFNα treatment, MDA5^946T^ SC-β cells displayed a higher number of unique DEGs when compared to their MDA5^627*^ counterpart, but the difference was only 1.4-fold. MDA5^627*^ SC-α and SC-δ cells exhibited 14.83-fold and 2.75-fold more DEGs, respectively, compared to their variant counterparts. Compared to the other cells within the same variant group, MDA5^946T^ SC-β cells displayed 10-fold or more the number of DEGs, while SC-α and -δ had similar numbers of unique genes (Fig. 3d). Within MDA5^627*^ cells, SC-α had the highest number of unique DEGs, with a little over two-fold and 11-fold more genes when compared to SC-β cells and -δ cells, respectively. SC-α and -δ cells harboring the MDA5^946T^ variant did not share any genes, while SC-α and -β cells from this same variant group shared 14 genes. SC-α and -δ cells from the MDA5^627*^ group shared 13 genes, while SC-α and -β cells shared 33 genes. SC-β and -δ cells across both groups shared the least number of genes (Fig. 3d).

CVB3-infected MDA5^946T^ cells shared 192 DEGs, whereas MDA5^627*^ cells shared 54 (Fig. 3e, Supplementary Data 7). MDA5^946T^ SC-β cells exhibited 6.71-fold more unique DEGs when compared to their MDA5^627*^ counterpart. MDA5^946T^ SC-α cells also displayed a strong unique transcriptional response to CVB3 infection that was only 1.15-fold lower than MDA5^946T^ SC-β cells. However, they only displayed 1.13-fold more unique genes than MDA5^627*^ SC-α cells. MDA5^946T^ SC-δ cells had the lowest number of unique DEGs, which was 18.09- and 15.78-fold lower compared to SC-β and -α cells from this variant group. MDA5^627*^-derived SC-α exibited the highest number of unique DEGs with 5.19- and 17.89-fold more when compared to MDA5^627*^ SC-β and -δ cells (Fig. 3e). MDA5^627*^ SC-δ cells had the lowest number of unique DEGs in this variant group and compared to MDA5^946T^ SC-δ cells. When looking at DEGs shared between paired cell types, MDA5^946T^ SC-α and -β cells shared 384 genes, while this cell type pair harboring MDA5^627*^ only shared 69 genes. SC-α and -δ cells shared 11 and 22 genes in the MDA5^946T^ and MDA5^627*^ groups, respectively. SC-β and -δ cells shared 46 and 10 genes in the MDA5^946T^ and MDA5^627*^ groups, respectively (Fig. 3e).

### Endocrine cell subpopulations exhibit distinct transcriptional signatures to different immune-associated stressors

To determine if the DEGs unique to each cell type were shared across stress conditions and variants, we first grouped all the genes by cell type and then graphed their log_2_ fold change. CVB3-infected MDA5^946T^ SC-α cells displayed a strong downregulation of multiple genes, including *ABCA2* (lipid transporter), *ELFN2* (phosphatase inhibitor), and *CLASRP* (regulates alternative splicing), compared to SC-α cells from all other conditions, including MDA5^627*^ cells (Fig. 3f, Supplementary Data 8). Infected MDA5^946T^ SC-α cells also upregulated multiple genes, including *ARRDC4* (a glucose transporter), *CMTM6* (PD-L1 regulator), *CYCS* (component of the mitochondrial electron transport chain [ETC] and pro-apoptotic gene), *FAM210B* (critical to mitochondrial function), *MPC2* (critical to mitochondrial function), and *ATP5MK* (subunit of mitochondrial ETC complex), that were downregulated in CVB3-infected MDA5^627*^ SC-α cells.

IFNα-treated SC-α cells from both variants upregulated multiple genes, including *SERPING1* (serine protease inhibitor), *ISG20* (IFN-induced gene), *PIK3KAP1* (chromatin regulator), *MOV10* (RNA helicase), and *CDK18* (stress response gene), and downregulated *ACO1* (regulates citric acid cycle), *NOP53* (regulates protein synthesis), and *MT-CYB* (ETC component) (Fig. 3f). However, we observed upregulation of multiple genes, such as *LATS1* (Hippo signaling component) and *BAK1* (pro-apoptotic gene), in IFNα-treated MDA5^627*^ SC-α cells only. Poly(I:C) treatment induced the downregulation of many ER- and mitochondria-associated genes (*SRP9*, *IARS2*, *SLC25A5*, *TIMM9*, *SCFD1*) and the upregulation of stress-associated genes (*ATF3*, *RASD1*, *FOS*, *ZFP36*) in MDA5^627*^ SC-α cells. In contrast, poly(I:C)-treated MDA5^946T^ SC-α cells induced differential expression of few genes compared to other treatments, with downregulation of *SRP9*, *LY6H*, *GTF2A2*, and *ECHS1*. Most DEGs in poly(I:C)-treated SC-α cells corresponded the MDA5^946T^ variant (Fig. 3f).

CVB3-infected MDA5^946T^ SC-β cells upregulated multiple genes, such as *KLF10* (involved in immune activation), *SSR1* (component of translocon, which helps move proteins across the ER), *TRMT10C* (a tRNA methyltransferase), and *SLC38A2* (an amino acid transporter), not found in other conditions or in MDA5^627*^ SC-β cells (Fig. 3g, Supplementary Data 8). Other genes unique to CVB3-infected MDA5^946T^ SC-β cells were associated with β cell development and function (*GLIS3*, *UCP2*, *RFX6*, *NKX6-1*), as well as the immune response (*DEXI*, *STAT6*, *TRIM41*), and were all downregulated. Infected MDA5^946T^ SC-β cells also upregulated *IFNAR1*, *GFPT1* (crucial for glycosylation), and *SSR3*, which were all downregulated in their MDA5^627*^ counterpart. CVB3 infection induced differential expression of unique genes in MDA5^627*^ SC-β cells, including the upregulation of genes involved in insulin secretion, metabolism, and protein processing (*RPN2*, *GIPR*, *ABBC8*, *MLXIPL*). In IFNα-treated SC-β cells from both variants, we observed upregulation of type I IFN-associated genes, including *ISG15* and *IFI6* (Fig. 3g). IFNα treatment also upregulated other immune response genes (*CXCL16*, *RFX5*, *IL10RB*) and signaling genes (*SMAD3*, *JUNB*) only in MDA5^946T^ SC-β. Following poly(I:C) treatment, *ISG15*, *HSPA1B*, and *IFI6*, which are all stress-associated genes, were only upregulated in MDA5^946T^ SC-β cells, while SC-β from both variants had lower expression of *BEX1* and *PRDX6*, both associated with mitochondrial stress (Fig. 3g).

*NLRC5*, a transcriptional activator of MHC/HLA class I genes, is highly expressed in IFNα-treated MDA5^946T^ SC-δ cells (Fig. 3h, Supplementary Data 8). IFNα also downregulated the expression of *MTURN* (NFκB signaling regulator), *ACACA* (involved in fatty acid synthesis), *GPC6* (involved in Wnt signaling), and *NUDT3* (regulates mRNA stability) in SC-δ cells from both variants, while *PRKAR2A* (regulates cyclic AMP signaling), *FIS1* (regulates mitochondrial fission), and *ERO1A* (regulates protein folding) were only lowly expressed in IFNα-treated MDA5^627*^ cells. *IFIT2* is highly expressed in both MDA5^946T^ and MDA5^627*^ SC-δ cells following IFNα treatment (Fig. 3h). CVB3 infection mostly decreased the expression of genes unique to MDA5^627*^ SC-δ cells. In MDA5^946T^ cells, CVB3 infection increased the expression of *SLC44A2*, *CLPX*, *MED10*, *PCDH1*, *DUSP6*, *ZNF709*, and *HSPA4L*, which are involved in processes such as protein folding and transcription. In CVB3-infected MDA5^946T^ SC-δ cells, we observed downregulation of *GP1BB*, *WNT4*, and *LRBA* (CTLA-4 regulator). *ACTG1*, an actin cytoskeleton component, expression decreased in all conditions and variants, while *SCLT1* was only upregulated in IFNα-treated cells. We also observed downregulation of genes involved in transcription, translation, and protein folding, including *MMGT1*, *TCEAL2*, *DPH3*, and *SLC29A1*, only in poly(I:C)-treated MDA5^627*^ SC-δ cells (Fig. 3h).

### Gene Ontology reveals cell- and variant-specific pathway enrichment

To understand the biological pathways affected by stress, we performed Gene Ontology (GO) analysis on all the DEGs presented in the Venn diagrams for each endocrine cell type and treatments across both variants. In SC-α cells, IFNα-enriched genes were associated with apoptosis and immune-related pathways in both variants (Fig. 4a-b, Supplementary Data 9). CVB3 infection in MDA5^946T^ SC-α cells led to the regulation of genes associated with the mitochondria, cell death, and the immune system. In contrast, we only observed associations with the ER and Golgi-associated vesicles in MDA5^627*^ SC-α cells following CVB3 infection. Poly(I:C)-enriched genes were linked to MAPK signaling and cellular response to stress in MDA5^627*^ cells, but no GO terms with a significant adjusted p-value (<0.05) were found in poly(I:C)-treated MDA5^946T^ SC-α cells (Fig. 4a-b).

**Figure 4.**
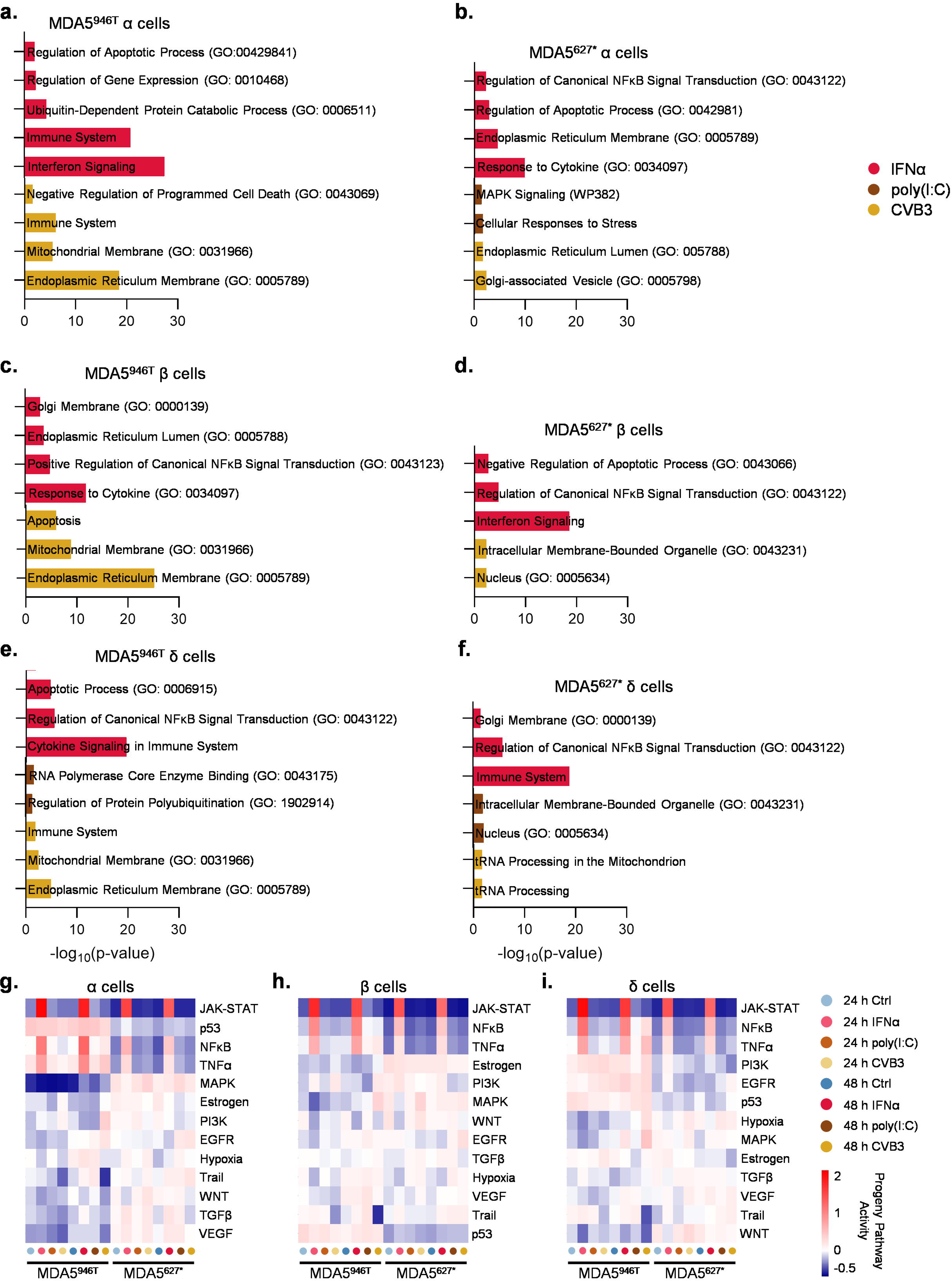
Gene ontology (GO) and PROGENy analyses of SC-islets reveal differences in cell signaling transcriptional signatures across cell types and MDA5 variants. **(A-F)**, GO analyses of pathways enriched in SC-α. -β, and -δ cells across both variants when comparing stress-treated cells to control. Time points were combined. **(G-I)** PROGENy analysis depicting the activity scores of 14 major pathways across time points and treatments in **(G)** SC-α, **(H)** SC-β, and **(I)** SC-δ cells (negative values = decreased activity; positive values = increased activity).

SC-β cells from both variants showed an enrichment of IFNα-associated genes linked to NFκB signaling and cytokine response (Fig. 4c-d, Supplementary Data 9). The transcriptional response to IFNα in MDA5^946T^ SC-β cells was also characterized by the enrichment of genes associated with the Golgi and ER, while apoptosis was highlighted in MDA5^627*^ cells. CVB3-enriched genes were linked to apoptosis, mitochondria, and the ER in MDA5^946T^ SC-β cells, whereas they were associated with the nucleus and intracellular membrane-bounded organelles in their MDA5^627*^ counterparts. We did not observe significant GO terms in poly(I:C)-treated SC-β cells from either variant.

A similar pattern of immune, Golgi, and apoptotic gene enrichment in response to IFNα was also observed in SC-δ cells (Fig. 4e-f, Supplementary Data 9). As with MDA5^946T^ SC-α and -β cells, CVB3 infection led to enrichment of genes linked to the immune system, mitochondria, and the ER. However, genes enriched in CVB3-infected MDA5^627*^ SC-δ cells were associated with tRNA processing and the mitochondria. In contrast, poly(I:C) treatment showed a different transcriptional profile, with associations to RNA polymerase and protein polyubiquitination in MDA5^946T^ cells (Fig. 4e). In MDA5^627*^ SC-δ cells, poly(I:C) treatment led to the differential regulation of genes associated with intracellular membrane-bounded organelles and the nucleus (Fig. 4f).

### PROGENy analysis uncovers distinct signaling pathway activity scores following stress

To determine the effects of each variant on signaling pathways following stress treatment, we utilized PROGENy^77^ to assess the directional activity of associated gene expression. In SC-α cells, MDA5^946T^ exhibited decreased activity scores for MAPK signaling and increases in p53 and TNFα, including at basal levels, when compared to MDA5^627*^ (Fig. 4g, Supplementary Data 10). We also noted differences in TGFβ activity scores between variants in all stress conditions except for CVB3 at 48 h. NFκB activity scores increased in both variants following IFNα treatment but were lower in MDA5^627*^ cells compared to MDA5^946T^ in all other conditions (Fig. 4g). JAK-STAT signatures were similar across both variants in these cells. PI3K signaling activity scores were high at 48 h of CVB3 infection in MDA5^946T^ SC-α cells but were low in MDA5^627*^ cells. Trail pathway activity was increased in stress-treated MDA5^627*^ cells (Fig. 4g).

In SC-β cells, TNFα signaling activity scores were increased following IFNα treatment in both variants, but they were only increased in MDA5^946T^ cells after 48 h of poly(I:C) and CVB3 treatment (Fig. 4h, Supplementary Data 10). NFκb, EGFR, TGFβ, and JAK-STAT activity score patterns were similar across all SC-β cells and resemble those of SC-α cells (Fig. 4g-h). MAPK values were low in MDA5^946T^ SC-β cells in all conditions, except for 48 h of CVB3 infection, while they are mostly high across all conditions with MDA5^627*^ cells. Similar to their SC-α cell counterpart, we observed high p53 activity scores for MDA5^946T^ SC-β cells across all conditions. MDA5^627*^ SC-β cells exhibited low Trail scores for the 24 h conditions when compared to SC-α cells of this group (Fig. 4g-h).

In SC-δ cells, activity score patterns for JAK-STAT, NFκB, and TGFβ resembled those of the other endocrine cells across variants (Fig. 4i, Supplementary Data 10). PI3K values were low in MDA5^627*^ SC-δ cells but showed opposite trends in their MDA5^946T^ counterparts. We observed TNFα increased following IFNα treatment in both variants, but its activity score was also increased following 48 h CVB3 infection in MDA5^946T^ cells (Fig. 4i). Unlike in the other endocrine cell types, PI3K and EGFR trended high in MDA5^946T^ SC-δ cells but low in MDA5^627*^ cells. Across both variants, Trail scores showed similar patterns to those of SC-β cells (Fig. 4i).

### Detailed transcriptional analysis of key signaling pathways reveal the MDA5^627*^ variant attenuates immune response in SC-islets

Given the shared association of multiple GO terms between SC-endocrine cell types, we investigated the specific gene signatures of these processes. NFκB signaling-associated genes were primarily linked to IFNα treatment. Our scRNA-seq analysis revealed strong similarities in gene expression across variants in all three cell types, with SC-α, -β, and -δ cells from both variants upregulating *BST2*, *STAT1*, *IFIT5*, *TRIM5*, *KLF4*, and *VEGFA* (Fig. 5a, Supplementary Data 11). However, a direct pairwise comparison of IFNα-treated variants revealed lower expression of *VEGFA* in all three MDA5^627*^ cell types, while *KLF4* and *BST2* expression were lower in MDA5^627*^ SC-α and -β, respectively (Fig. 5a). MDA5^946T^ SC-α cells also had higher expression of *TICAM1*, *TFRC*, *MTDH*, *SQSTM1*, *TNFSF10*, *MYD88*, and *CD74* compared to MDA5^627*^ counterparts. Although *TICAM1*, *MTDH*, and *MYD88* expression was also higher in MDA5^946T^ SC-β cells compared to their variant counterpart, *TNFSF10* expression was lower, with higher expression of *SHISA5* and *DAB2IP* (Fig. 5a). MDA5^946T^ SC-δ cells exhibited higher expression of *MTDH*, *SQSTM1*, *MYD88*, and *DAB2IP* compared to MDA5^627*^ SC-δ cells.

**Figure 5.**
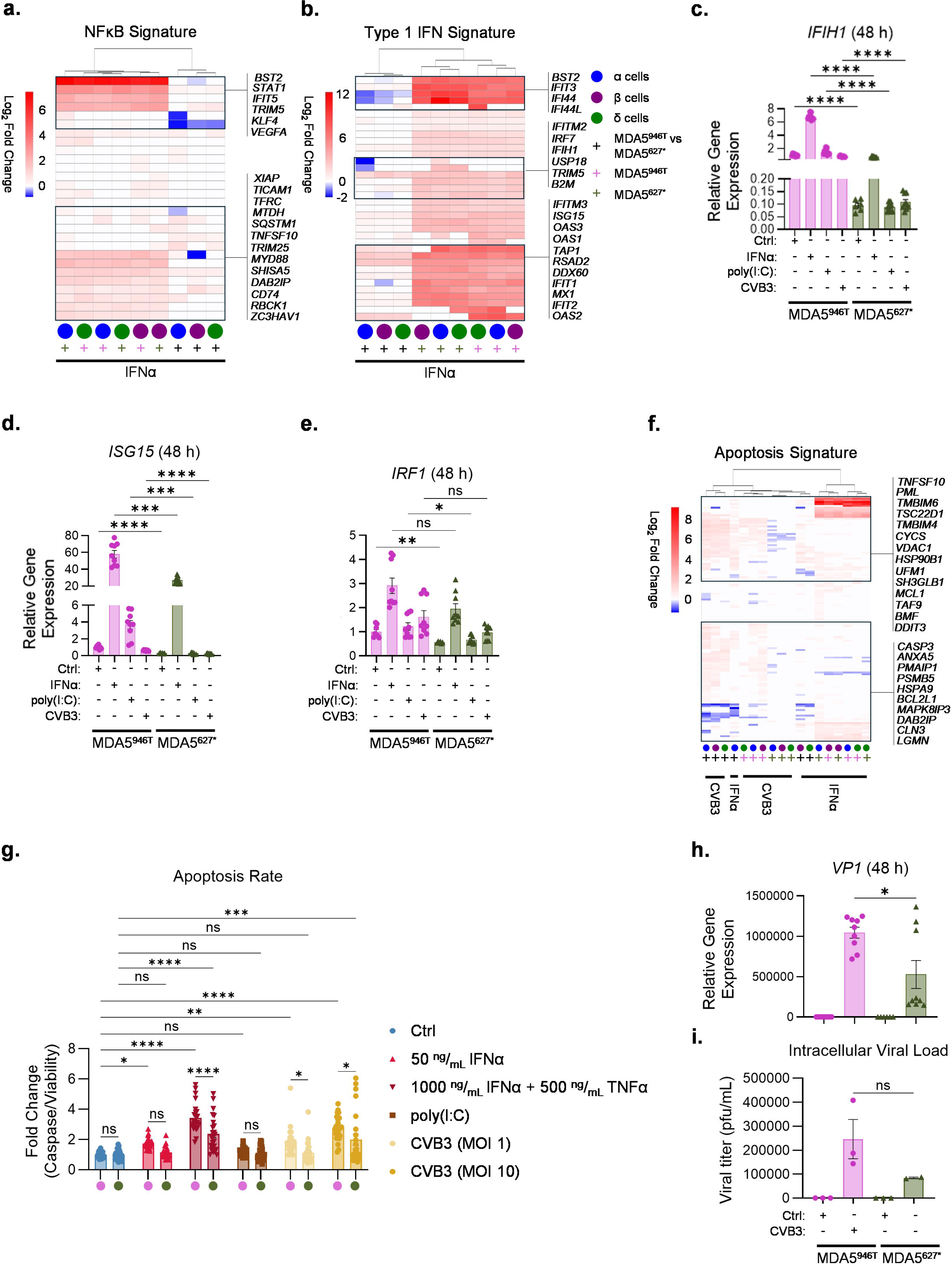
scRNA-seq reveals a dampened response to stress treatment in MDA5^627*^ SC-islets. **(A-B)**, Heatmaps depicting log_2_ fold change of **(A)** NFκB signaling-associated and **(B)** type 1 IFN-associated DEGs across SC-α, -β, and -δ cells when comparing the corresponding IFNα treatment to control or stress-treated MDA5 variants to each other. Positive fold change values (red) in variant-to-variant comparisons correspond to upregulation in MDA5^946T^ cells when compared to their MDA5^627*^ counterparts. **(C-E)**, rt-qPCR of type 1 IFN-associated genes (n = 3). Error bars represent s.e.m.** = Brown-Forsythe and Welch ANOVA tests followed by Dunnett’s T3 multiple comparisons test. **(F)**, Heatmap depicting log_2_ fold chance of apoptosis-associated DEGs across SC-α, -β, and -δ cells when comparing the corresponding stress treatment to control or stress-treated MDA5 variants to each other. Positive fold change values (red) in variant-to-variant comparisons correspond to upregulation in MDA5^946T^ cells when compared to their MDA5^627*^ counterparts. **(G)**, Apoptosis assay measuring caspase 3/7 activity normalized to cell viability (n = 3). Error bars represent s.e.m. ** = Ordinary two-way ANOVA followed by Sidak’s multiple comparisons test. **(H)**, rt-qPCR of viral genome expression in CVB3-infected SC-islets (n = 3). Error bars represent s.e.m. ** = Unpaired t test. **(I)**, Intracellular viral titer in SC-islets following 48 h of CVB3 infection (n = 3). Error bars represent s.e.m. ** = Welch’s unpaired t test. Outlier identified in infected MDA5^627*^ SC-islet samples using the Grubbs’ method.

As expected, IFNα treatment led to the strong upregulation of many type I IFN-associated genes (Fig. 5b, Supplementary Data 11). Most of the investigated genes were upregulated in SC-endocrine cells from both variants. All three MDA5^946T^ cell types showed higher expression of these genes compared to MDA5^627*^ cells, with some exceptions (Fig. 5b). *BST2*, *B2M*, and *RSAD2* expression was lower in MDA5^946T^ SC-β cells compared to their MDA5^627*^ counterpart, while *IFITM2* and *IRF7* were both lowly expressed in MDA5^946T^ SC-α cells compared to MDA5^627*^ SC-α cells. All three MDA5^946T^ SC-endocrine cell types had lower expression of *IFI44* compared to their variant counterparts, with *IFI44L* expression also being lower in MDA5^946T^ SC-α and -β cells (Fig. 5b).

We validated the observations from our scRNA-seq data using whole SC-islets treated with all stressors and running real-time quantitative PCR (rt-qPCR). Basal levels of *IFIH1*, *ISG15*, and *IRF1* were significantly lower in MDA5^627*^ SC-islets, with a nonsignificant trend in *IRF1* at 24 h (Fig. 5c-e, Extended Data Fig. 5a-c, Supplementary Data 2). We observed increases in *IFIH1*, *ISG15*, and *IRF1* following IFNα treatment for 24 and 48 h, with little to no increase in poly(I:C)- and CVB3-treated SC-islets (Fig. 5c-e, Extended Data Fig. 5a-c). MDA5^627*^ SC-islets had significantly lower gene expression for these genes across all treatments. The exception was *IRF1* at 24 h for all treatments and at 48 h for IFNα only. *DDX58*, which encodes the RNA helicase retinoic acid-inducible gene I (RIG-I), was also lower in MDA5^627*^ SC-islets across all conditions except for IFNα treatment at both time points (Extended Data Fig. 5d-e). *DDIT3*, a pro-apoptotic marker, was basally higher in MDA5^627*^ SC-islets but did not increase with treatment (Extended Data Fig. 5f-g). *IFIT2* did increase with IFNα treatment in both variants with no significant differences (Extended Data Fig. 5h-i).

### MDA5^627*^ reduces apoptosis and viral titer in SC-islets

Apoptosis- and cell death-associated GO terms were also significantly enriched in CVB3-and IFNα-treated cells (Fig. 4a-f). Therefore, we investigated the expression of associated genes in SC-endocrine cells across both variants. All CVB3-infected MDA5^946T^ SC-endocrine cells had an overall stronger apoptotic transcriptional signature compared to their direct MDA^627*^ counterparts, with higher expression of multiple genes such as *TMBIM6*, *CYCS*, *SH3GLB1*, *ANX5* and *PMAIP1* (Fig. 5f, Supplementary Data 11). Moreover, MDA5^946T^ SC-β cells had the most DEGs within its variant group, with mostly upregulated genes. CVB3-infected MDA5^627*^SC-endocrine cells differentially regulated few apoptosis-associated genes, with downregulation of *HSP10B1* and *UFM1*. Additionally, there was little overlap between DEGs induced by CVB3 infection and TNFα treatment. *PML*, *DAB2IP*, *CLN3*, and *LGMN* were all enriched in TNFα-treated cells from both variants (Fig. 5f). IFNα-treated MDA5^627*^ SC-α cells exhibited unique upregulation of several genes, including *ATF4* and *RHOB*, while their MDA5^946T^ counterparts did not differentially express many unique genes (Supplementary Data).

To determine if our apoptotic transcriptional data translated to changes in apoptotic rates, we multiplexed the CellTiter-Fluor Cell Viability Assay with the Caspase-Glo 3/7 Assay System on plated down SC-islets from both variants. IFNα treatment alone significantly increased apoptosis in MDA5^946T^ SC-islets compared to control but not in MDA5^627*^ cells, with a significant difference when directly comparing the variants (Fig. 5g). Since we did not observe significant cell death of MDA5^627*^ SC-islets with IFNα treatment alone, we cultured SC-islets from both variants with a cytokine mixture composed of IFNα and TNFα (Fig. 5g). This mixture induced apoptosis in SC-islets from both variants, with significantly higher apoptotic rates in MDA5^946T^ SC-islets compared to MDA5^627*^ (Fig. 5g). As expected, we did not see significant changes in apoptosis following poly(I:C) treatment compared to control in both groups. CVB3 infection with a MOI of 1 significantly increased apoptosis in MDA5^946T^ SC-islets compared to control SC-islets but not in MDA5^627*^ cells. A direct comparison between the two variants to each other revealed a significant difference. Increasing the MOI to 10 significantly increased apoptotic rates in both variants compared to control groups, with significantly higher rates in MDA5^946T^ SC-islets compared to MDA5^627*^ SC-islets (Fig. 5g).

Given the lower immune response and apoptotic rates in MDA5^627*^ SC-islets, we also measured infection rates in both variants. MDA5^627*^ SC-islets had significantly lower viral genome expression when compared to MDA5^946T^ SC-islets at both time points (Fig. 5h, Extended Data Fig. 5j, Supplementary Data 2). Plaque assays measuring intracellular virion levels revealed a similar trend, though it was not a significant difference (Fig. 5i).

### MDA5^627*^ dampens stress-mediated mitochondrial dysfunction and impaired islet function

Building upon our GO analysis, which revealed a strong association with mitochondrial-related terms in CVB3-infected cells, we investigated both mitochondrial gene signatures and function. Across variants, CVB3 infection had a stronger transcriptional impact on MDA5^946T^ SC-α, -β, and -δ cells, leading to the upregulation of several genes unique to this variant, including *VDAC1*, *SLC25A33*, *COX6C*, and *NDUFAB1*, when compared to untreated control cells (Fig. 6a, Supplementary Data 11). In contrast, MDA5^627*^ cells showed only a mild mitochondrial-associated transcriptional response to CVB3 infection, with most DEGs being downregulated. MDA5^627*^ SC-α cells exhibited the highest number of DEGs of all cell types in this variant, with decreased expression of *ATP5MK*, *CYCS*, *SLC25A33, COX6C*, and *NDUFB4* (Fig. 6a). While MDA5^946T^ SC-β cells showed the strongest transcriptional response within its variant group, we only observed four downregulated DEGs in MDA5^627*^ SC-β cells compared to their respective untreated controls. A direct comparison between variants confirmed that all three MDA5^946T^ cell types had higher expression of most genes investigated, although *MT-CO1* and *MT-CO2* were both downregulated when compared to all MDA5^627*^ cell types (Fig. 6a).

**Figure 6.**
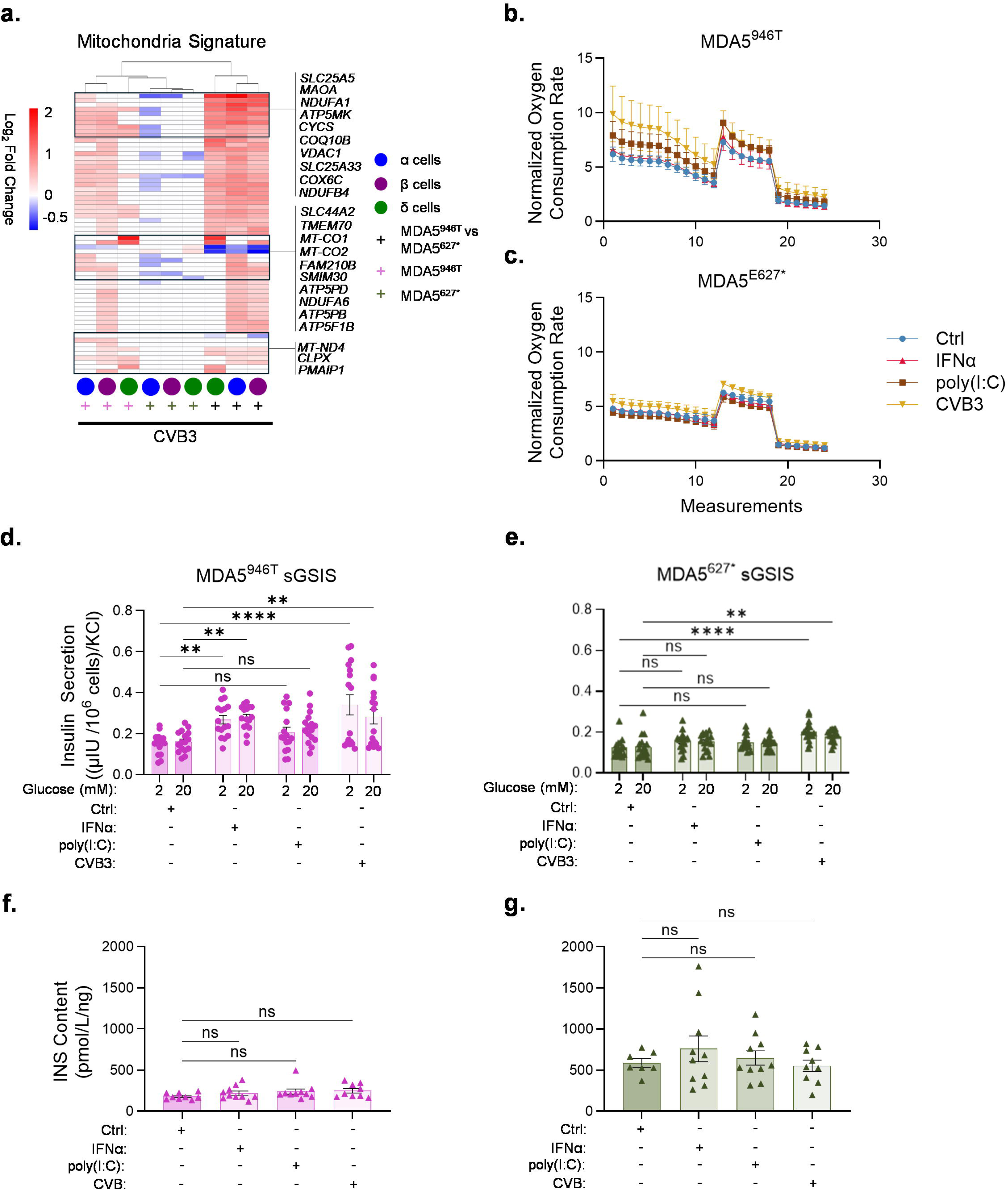
The MDA5^627*^ variant dampens stress-mediated mitochondrial and insulin secretion dysfunction in SC-islets. **(A)**, Heatmap depicting log_2_ fold chance of mitochondria-associated DEGs across SC-α, -β, and -δ cells when comparing CVB3-infected cells to control or stress-treated MDA5 variants to each other. Positive fold change values (red) in variant-to-variant comparisons correspond to upregulation in MDA5^946T^ cells when compared to their MDA5^627*^ counterparts. **(B-C)**, Plots showing normalized mitochondrial oxygen consumption rate in SC-islets following 48 h stress treatment before and after injection of Oligomycin, FCCP, and Antimycin A/Rotenone (n = 2). Data normalized to DNA content. Error bars represent s.e.m. **(D-E)**, Statis glucose-stimulated insulin secretion of SC-islets treated with various stressors for 48 h. Normalized to KCl-stimulated insulin secretion (n = 4). Error bars represent s.e.m. ** = RM Two-way ANOVA followed by Dunnett’s multiple comparisons test. **(F-G)**, Intracellular insulin content measurements following SC-islet treatment with various stressors for 48 h (n = 3). Error bars represent s.e.m. ** = Ordinary one-way ANOVA with Dunnett’s multiple comparisons test.

To determine if these transcriptional changes impacted cellular metabolism, we assessed mitochondrial function by measuring oxygen consumption rates (OCRs) in plated down SC-islets using the Seahorse XF system. We also included other treatments to further validate our findings. IFNα-treatment did not induce any changes in OCRs across all measurements in MDA5^946T^ SC-islets (Fig. 6b). Poly(I:C)-treated MDA5^946T^ SC-islets had increased respiration at basal levels and following oligomycin and carbonyl cyanide-4 (trifluoromethoxy) phenylhydrazone (FCCP) injections. CVB3 infection induced the biggest effect in these cells, with increased OCRs throughout the entire assay (Fig. 6b). In contrast, MDA5^627*^ SC-islets did not have any changes in OCRs when treated with IFNα or poly(I:C), with only a mild increase in OCR following oligomycin and FCCP injections of CVB3-infected SC-islets (Fig. 6c).

Given the role of mitochondria in pancreatic islet function and that SC-islet function and identity can be negatively affected by stress,^39^ we investigated whether the MDA5^627*^ allele could help preserve islet function. We treated SC-islets with the various stressors for 48 h. sGSIS on MDA5^946T^ SC-islets revealed that poly(I:C) had no effect on insulin secretion (Fig. 6d). However, both IFNα and CVB3 treatment significantly increased insulin secretion. In contrast, MDA5^627*^ SC-islets significantly increased insulin secretion only when treated with CVB3 (Fig. 6e). Despite these increases in secretion, insulin content and *INS* gene expression remained unchanged following treatment (Fig. 6f-g, Extended Data Fig. 6a-b). Additionally, gene expression of other islet hormones did not change following treatment for 24 and 48 h (Extended Data Fig. 6c-f), except for increases in *GCG* within IFNα- and CVB3-treated MDA5^946T^ SC-islets and reductions in *SST* expression following CVB3 infection in this same variant at 48 h (Extended Data Fig. 6d, f). SC-islets from both variants did exhibit reductions in *NKX6-1* expression with 48 h CVB3 infection, despite no changes in *CHGA* expression (Extended Data Fig. 6g-j).

## Discussion

GWAS studies have identified *IFIH1* as a T1D-associated gene, with both risk and protective SNPs characterized in PBMCs.^32–34,36,37^ Others have shown that *Ifih1* deficiency in mice delayed T1D onset and reduced islet inflammation following CVB3 and CVB4 infection.^70,78^ These studies suggest that a dampened MDA5-mediated immune response is protective. However, the association between *IFIH1*, immune response, islet health and function, and T1D had not been investigated in a human pancreatic context. Such studies are difficult to conduct due to the scarcity of isolated islets from affected patients and that any isolated cells are of low quality. Here, we utilized CRISPR-Cas9-edited hPSC-derived pancreatic islets from a T1D patient to identify the effects of the protective MDA5^627*^ allele on transcriptional and functional responses at both a whole islet and single-cell resolution. Our SC-islets contained all relevant pancreatic endocrine cell types and were capable of secreting insulin in response to glucose stimulation. MDA5 protein expression increased in MDA5^946T^ SC-islets following treatment with high concentrations of IFNα but was undetectable in both control and treated MDA5^627*^ SC-islets. The failure to detect the truncated MDA5 protein via western blot suggests a major issue with protein stability and/or folding. This absence is compounded by a reduction in basal *IFIH1* mRNA levels, which limits the translation pool. Future studies should investigate the protein structure of the truncated MDA5 to determine if the C-terminal truncation induces severe protein misfolding, resulting in either masking of the Arg470 epitope or rapid, constitutive protein degradation.

We found that following stress treatment, MDA5^946T^ SC-islets exhibited a stronger transcriptional response to IFNα and CVB3—but not poly(I:C)—compared to MDA5^627*^ SC-islets. Furthermore, we discovered cell-type specific transcriptional responses to each stressor. Our functional data validated these transcriptional differences, revealing that the MDA5^627*^ variant favorably impacted apoptosis, mitochondrial function, viral replication, and insulin secretion.

Previous studies in PBMCs, cell lines, and mice have suggested that MDA5^946T^ leads to an increased basal type I IFN signature and a stronger immune response to stressors.^33–35^ Our findings in human SC-islets corroborate these observations. Pairwise comparisons at baseline revealed that MDA5^946T^ SC-islets exhibited higher expression of type I IFN-associated genes, which we confirmed by rt-qPCR showing lower basal *IFIH1*, *ISG15*, and *IRF1* gene expression in MDA5^627*^ SC-islets. This elevated basal activity in the MDA5^946T^ variant has been previously attributed to increased ATPase activity in MDA5.^34^ It may also be caused by increased response to endogenous dsRNA.^33^ In contrast, the MDA5^627*^ variant has been reported to lead to reductions in *IFIH1* gene expression, a finding that is consistent with our observations, and a complete loss of dsRNA binding activity.^36,38^ It also reduced type I IFN response to IFNβ and poly(I:C) treatment in PBMCs.^36,37^ While we do not make claims regarding the functional and regulatory changes that impact MDA5 within SC-islets, these molecular deficiencies provide a clear mechanistic explanation for the dampened immune response and reduced apoptosis we observed.

Building on these basal differences, our pseudo-bulk and single cell analyses further revealed that MDA5^946T^ cells underwent a stronger transcriptional response, marked by a higher number of DEGs, to IFNα and CVB3 treatment compared to MDA5^627*^ cells. This pattern was not observed following poly(I:C) treatment, suggesting the MDA5^627*^ SC-islets may activate alternative, non-canonical pathways to compensate for the lack of dsRNA activity. Our PROGENy analyses also showed distinct patterns in how the variants impact cellular signaling pathways in response to stress, with more pronounced increases in activity scores of pathways such as p53 and TNFα in MDA5^946T^ cells. We also showed that type I IFN- and NFκB-associated gene expression was consistently higher in MDA5^946T^ SC-α, -β, and -δ cells compared to their MDA5^627*^ counterparts. Moreover, the higher expression of *TAP1* and *TAP2*, which are involved in antigen presentation to the immune system, in IFNα-treated MDA5^946T^ SC-islets suggests a potential mechanism by which the MDA5^627*^ variant may also protect against T1D. As the islets of autoantibody-positive and T1D patients hyper-express class I HLA molecules, it is possible the MDA5^627*^ variant leads to decreased autoantigen presentation to immune cells, a critical step for T cell infiltration into the pancreas.^79–82^

A recent study reported that CVB3 infection induces unique, cell type-specific transcriptional responses in human primary islets, with strong impacts on β and α cells.^67^ Our data from SC-islets are consistent with these findings. Here, we also report that SC-β and SC-α cells were the most transcriptionally impacted pancreatic endocrine cell types, not only to CVB3 infection but also to IFNα treatment. The muted response to poly(I:C) has also been previously reported in pancreatic islets.^67^ However, our findings reveal that this stress response is highly variant-dependent. MDA5^946T^ SC-islets had the strongest transcriptional response in SC-β cells, whereas MDA5^627*^ SC-islets showed the strongest response in SC-α cells. For instance, 48 hours of CVB3 infection induced 897, 762, and 371 DEGs in MDA5^946T^ SC-β, -α, and -δ cells, respectively, while it induced 222, 373, and 128 DEGs in MDA5^627*^ SC-β, -α, and -δ cells, respectively. Moreover, we found a distinct pro-apoptotic signature in the MDA5^946T^ variant that corroborates and expands on our observations. Following CVB3 infection, MDA5^946T^ SC-β and - α cells had a greater number of upregulated apoptosis-associated genes compared to MDA5^627*^ cells, with SC-β cells being the most sensitive. For instance, in MDA5^946T^ SC-β cells, pro-apoptotic genes, such as *CASP3*, *CYCS*, and *PMAIP1*, were highly upregulated following CVB3 infection, while in their MDA5^627*^ counterpart, these genes were not significantly changed. These observations directly explain the significantly higher apoptotic rates we measured in these islets following IFNα and CVB3 treatment. They also highlight a mechanism by which the MDA5^627*^ variant confers protection, not just by dampening a broad immune response, but also by specifically reducing the pro-apoptotic signature that is highly detrimental to β cells.

Viral infections have been shown to impact the mitochondria to favor virion generation.^83,84^ In primary human islets, CVB3 infection induced differential expression of mitochondria-associated genes, mitochondrial dysfunction, and differential effects on mitochondrial size in β and α cells.^67^ Our findings also support some of these claims. GO analyses revealed enrichment of mitochondria-associated terms in CVB3 infected cells. MDA5^946T^ SC-islets had strong induction of several mitochondria-associated genes, including *SLC44A2*, *TMEM70*, *MT-ND4* and *CLPX*, that were not differentially expressed in MDA5^627*^ cells. In fact, we observed mostly downregulated mitochondrial genes in MDA5^627*^ SC-endocrine cells. Our functional data also showed mitochondrial dysfunction only in MDA5^946T^ SC-islets following CVB3 infection. In contrast, mitochondrial function was maintained in MDA5^627*^ SC-endocrine cells. This may be caused, in part, by the lower levels of viral genome and titer we observed in MDA5^627*^ SC-islets, as decreased viral levels would result in lower impacts on mitochondrial function^83,84^. These benefits to mitochondrial function may also help further promote β cell survival and function during stress, given that β cells are highly susceptible to mitochondrial stress and that insulin secretion requires high respiration rates.^85–87^ Our data showing decreased impacts on insulin secretion following stress in MDA5^627*^ SC-islets corroborate this notion. The MDA5^627*^ variant may also dampen negative effects on insulin secretion by simply preserving β cell identity and promoting cell survival, as we have shown. Together, these findings suggest that the MDA5^627*^ variant confers protection against T1D by preventing viral- and inflammatory-induced mitochondrial dysfunction, thereby preserving islet metabolic health and function.

This study has some limitations that can be addressed in future studies. First, we investigated the impacts of the protective MDA5^627*^ variant in a T1D genetic background using hPSCs derived from only one T1D patient. To further validate the generalizability of our findings and capture the genetic heterogeneity found in the population, our dataset could be expanded to include SC-islets generated from other hPSC lines, including those from other T1D and genetically susceptible non-diabetic patients. Cross-referencing our dataset with previous work using scRNA-seq to study primary islets and SC-islets under various stressors, including cytokines, CVB3/4, and small molecules, could provide new insights.^47,66,67,88–96^ Future work should also investigate how MDA5^627*^ protects against CD8+ T cell-mediated β cell destruction by using T cells derived from isogenic lines. Another limitation is the presence of the off-target EC populations. The lower basal function of MDA5^946T^ SC-islets could be attributed to the higher proportion of EC cells, which have been reported to negatively affect insulin secretion.^47,97^ Considering this, future studies should explore whether the MDA5^627*^ improves SC-islet differentiation outcome by reducing SC-EC cell generation. Furthermore, given that SC-islets are functionally and transcriptionally immature, their use may provide an incomplete understanding of the effects of MDA5^627*^ on islet health and function in response to stress.^92–94,98^ To more accurately model the chronic nature of T1D and determine if the observed protective effects of the MDA5^627*^ variant are sustained over time, further work is needed using longer time points.

In the present study, we demonstrated the utility of our dataset in performing functional studies to characterize the role of genetic variants in SC-islet function and cell identity. We are the first to report that MDA5^627*^ leads to a dampened immune response and cell type-specific transcriptional differences compared to a full length MDA5 protein, as well as protects β cells against stress-induced dysfunction and cell death, advancing the understanding of the link between genetic background and T1D pathogenesis. Our study further validates SC-islets as a robust platform for studying clinically relevant genetic variants and their functional consequences in a human context. Overall, our work highlights variant-specific differences in response to stress treatments, providing insights that would improve preventative therapies against T1D.

## Methods

### Approvals

All research was conducted in compliance with relevant ethical guidelines. The design of the study did not incorporate sex as a variable, as this was beyond the scope of the investigation. This research received approval from the Washington University Institutional Biological & Chemical (IBC) Safety Committee (Approval number 20889).

### CH1-064 culture

The human induced pluripotent stem cell (hPSC) line, CH1-064, was cultured in mTeSR1 medium (Stem Cell Technologies, 05850) at 37°C with 5% CO_2_. Cells were passaged every 3-4 days. Briefly, cultures were washed with 1X PBS (Corning, 21-030-CV) and then incubated with TrypLE Express (Thermo Fisher, 12604-039) at a volume of 0.2 mL/cm^2^ for up to 7 minutes at 37°C. To neutralize the enzyme, an equal volume of mTeSR1 supplemented with 10 μM Y-27632 27632 (Pepro Tech, 129382310MG) was added. Following dissociation, cells were counted using a Vi-Cell XR (Beckman Coulter) and centrifuged at 300 x *g* for 3 min. The supernatant was discarded, and the cell pellet was resuspended in supplemented mTeSR1. Cells were then plated onto Matrigel-coated plates at a density of 0.8 x 10^5^ cells/cm^2^ for propagation. The culture medium was supplemented with Y-27632 for the initial 24 h to improve cell survival, after which daily media changes were performed using mTeSR without the supplement.

### CRISPR-Cas9 gene editing of *IFIH1* at *rs1990760* and *rs35744605*

CH1-064 hPSCs at ∼70% confluency were dissociated to single cells with Accutase (Stem Cell Technologies, 07920) and edited to carry either the *IFIH1 rs1990760^T/T^* (MDA5^946T^), or *rs35744605* (MDA5^627*^) variants using CRISPR-Cas9 technology. Gene editing was performed by homology-directed repair (HDR) with a double-stranded DNA repair template containing homology arms flanking the *IFIH1* target region. Guide RNAs were designed in CRISPOR (UC Santa Cruz), selected for high predicted specificity and distance of the Cas9 cut site to the single nucleotide polymorphism (SNP), and synthesized by Synthego. For each SNP, the HDR template introduced the desired base substitution, a synonymous change that removed a TspR1 restriction site for screening, and a disruption of the closest downstream protospacer adjacent motif (PAM) site to either *rs1990760* or *rs35744605*. Single guide RNA (7.5 µM) targeting *IFIH1* (*rs1990760* or *rs35744605*) was complexed with Cas9 protein (1.5 µM, Invitrogen, A36498). Cells were resuspended in P3 buffer (Lonza, V4XP-3032) and nucleofected with the sgRNA–Cas9 complex using program #CA-137 on the 4D-Nucleofector X Unit (Lonza). Transfected cells were plated on Matrigel-coated (Corning, 354277) 12-well plates in 1 mL mTeSR Plus medium (Stem Cell Technologies, 05825) supplemented with CloneR (Stem Cell Technologies, 05888), 10 µM Y-27632 (Stem Cell Technologies, 72302), and 2 µM NU7441 (Tocris, 3712). Cells were cultured for 4 days with daily medium changes until they reached ∼70% confluency.

At ∼70% confluency, hPSCs were dissociated to single cells with Accutase. Approximately 15,000 cells were replated on a Matrigel-coated 10-cm dish in 10 mL mTeSR Plus supplemented with CloneR for clonal isolation. The remaining cells were pelleted, and genomic DNA was extracted using Lucigen Quick Extract (Sigma, LGCQE09050) to assess bulk editing by Sanger sequencing. Chromatograms were analyzed with EditR to estimate editing frequency. Medium on clonal plates was changed every other day until individual colonies were large enough to pick. Single colonies were expanded and screened by genomic DNA extraction followed by Sanger sequencing to confirm correct editing. Clones that were homozygous for each variant were expanded further and evaluated for normal karyotype and retention of pluripotency marker expression via flow cytometry using an Accuri flow cytometer. Data was analyzed on FlowJo version 10.6.

### Western blot

For protein analysis, immunoprecipitations were performed using Dynabeads Protein A Immunoprecipitation kit (Thermo Fisher, 10006D). Briefly, cells were incubated on ice for 5 min in lysis buffer (Cell Signaling, 9803) supplemented with 1mM phenylmethylsulphonyl fluoride. Cells were sonicated for 3 sec 3x on ice with a probe sonicator (Sonics) and centrifuged for 10 min at 14,000 x *g* at 4°C. The supernatant was collected, and protein concentration was determined by BCA (Thermo Fisher, 23227). Dynabeads were linked to MDA5 antibody (Cell Signaling, 5321), which was diluted at 1:100, and rotated for 10 min at room temperature. 500μg protein was added to MDA5 bound beads and rotated overnight at 4°C. The following day, beads were washed, and protein was eluted and separated on an 8% Bis-Tris gel (Thermo Fisher, WG1002A) in MOPS running buffer (Thermo Fisher, B0001). The proteins were transferred onto PVDF membrane (Thermo Fisher, PI88520), blocked in 5% milk for 1 h at room temperature, and incubated in primary antibody in 5% BSA/TBS overnight at 4°C. Primary antibody, MDA5, was diluted at 1:1000. The following day, the blot was washed and incubated in Clean blot IP detection reagent (Thermo Fisher, 21230) in 5% milk for 1 h at room temperature before being detected with Cytiva ECL reagent (Amersham, RPN3004) on an Odyssey FC (Licor). For the Clean blot, the secondary antibody was used at 1:1000.

### CH1-064 differentiation

CH1-064 hPSCs were differentiated into pancreatic stem cell-derived islets (SC-islets) following our recent protocol.^68^ Initially, hPSCs were dissociated into a single-cell suspension and seeded at a density of 0.66 x 10^6^ cells/cm^2^ in mTSeR1 medium supplemented with Y-276352. After 24 h, the medium was replaced with 0.33 mL/cm^2^ of a differentiation base medium containing the appropriate factors for the first stage of differentiation. For MDA5^946T^ hPSCs, the final latrunculin A (LATA) concentration for this first medium change was 0.075 μM while it was 0.130μM for MDA5^627*^ hPSCs. Subsequent daily media changes were made with 0.33 mL/cm^2^ of differentiation base medium containing the corresponding factors. On day 1 of stage 5, we used a final 1.5 μM LATA concentration for both variants. On day 1 of stage 6, the cells were again dissociated into a single-cell suspension and re-plated at a density of 10 x 10^6^ cells per well in a 6-well plate with 5 mL of the corresponding differentiation media. The plates were then placed on an Orbi-Shaker set to 100 RPM to promote the formation of 3D cell aggregates. Daily media changes were performed for the first 48 h post aggregation. Media changes were performed every other day afterwards. The SC-islet clusters were harvested for downstream assays between day 18 and 21 of stage 6 of the differentiation protocol.

### Flow cytometry

Samples were single-cell dispersed with TrypLE Express for 15 min at 37°C, resuspended with an equal volume of 1X PBS and centrifuged at 300 x *g* for 3 min. After removing the supernatant, 4% paraformaldehyde (Electron Microscopy Sciences, #157-4-100) was added to the tubes, which were immediately mixed vigorously to disperse the cell pellet. After fixing for 30 min at 4°C, the cells were washed once with 1X PBS. Prior to processing for flow cytometry, samples were stored in 1X PBS and kept in 4°C. Dispersed cells were incubated in ICC solution (PBS + 0.1% Triton-X + 5% donkey serum) for 45 min on ice. Primary antibody solutions were made in ICC solution as follows: 1:300 rat anti-C-peptide (DSHB, GN-ID4-S), 1:1000 mouse anti-NKX6.1 (DSHB, F55A12), 1:1000 rabbit anti-CHGA (Abcam, ab15160), 1:250 anti-SST (BD, 566032), 1:350 anti-GCG (BD, 565891) and 1:500 anti-SLC18A1 (Sigma, HPA06379). Secondary antibodies were diluted 1:500 in ICC solution as follows: anti-rat PE (Jackson ImmunoResearch, #712-116-153), anti-mouse Alexa Fluor 488 (Invitrogen, A21202) and anti-rabbit Alexa Fluor 647 (Invitrogen, A31573).

### Static glucose stimulated insulin secretion assay

Glucose-stimulated insulin secretion (GSIS) was performed on SC-islets as previously described.^42^ Briefly, SC-islets were washed three times with Krebs-Ringer bicarbonate (KREB) buffer. They were then pre-incubated for one hour at 37°C in KREB buffer containing 2 mM glucose. This was followed by sequential one-hour incubations in KREB buffer with a low concentration of glucose (2 mM), a high concentration of glucose (20 mM), and then 30 mM KCl. The insulin concentration in the supernatant from each incubation was measured using a human insulin-specific ELISA kit (ALPCO, #80-INSHU-E01.1) according to the manufacturer’s instructions.

### Insulin content measurement

To measure the hormone content, whole SC-islet clusters were first washed twice with 1X PBS. The clusters were then incubated with a solution of 1.5% HCl in 70% ethanol in Eppendorf tubes. These tubes were stored at -20°C for 72 h and vortexed vigorously every 24 h to ensure complete insulin extraction. Following extraction, samples were centrifuged at 2,100 RCF for 15 min. The supernatant from each sample was collected and neutralized with an equal volume of 1 M Tris (pH 7.5). The concentrations of insulin were then quantified using a human insulin-specific ELISA kit. To normalize the data, the final insulin content was expressed relative to total DNA content, which was quantified using the Quant-iT Picogreen dsDNA Assay Kit (ThermoFisher, P7589) according to the manufacturer’s instructions.

### SC-islet infection and stress treatment

On treatment day, one well containing SC-islets was single-cell dispersed by incubating in TrypLE Express for 15 min. The single-cell solution was counted using a Vi-Cell XR. These cell counts were used to determine the amount of CVB3-Woodruff (titer = 1.98 x 10^12^ pfu/mL) to add to a new well containing an equal amount of SC-islet clusters. Whole SC-islets were then treated with either endotoxin-free water (control), 50 ng/mL IFNα (R&D Systems, 10984-IF), 500 ng/mL poly(I:C) (InvivoGen, tlrl-piclv), or CVB3-Woodruff (titer = 1.98 x 10^12^ pfu/mL) at a MOI of 20. Cells were collected for downstream assays at 24 and 48 h post-treatment.

### Cell preparation for scRNA-seq

Single-cell transcriptomic profiling was performed using the Chromium Single Cell Fixed RNA Profiling v1 Kit (10X Genomics, #1000414, #1000476) according to the manufacturer’s protocol. Briefly, SC-islets were dissociated into a single-cell suspension by incubating them with TrypLE Express for 15 min at 37°C. The resulting single-cell suspensions were then fixed and hybridized with multiplexing probes following the 10X Genomics Fixed RNA Profiling protocols (CG000478, CG000527).

### scRNA-seq

The hybridized samples were submitted to the Washington University McDonnell Genome Institute for library preparation and sequencing. Libraries were constructed using the Chromium Single Cell 3’ v3.1 Library and Gel Bead Kit (10X Genomics). The resulting libraries were then sequenced on an Illumina NovaSeq6000 System.

### scRNA-seq analysis

All single-cell analyses and comparisons were performed using Seurat v5.3.0 in R Studio (R version 4.4.0).^99^

### Raw data processing

The sequenced reads were aligned to the human GRCh38 reference genome, and the resulting data underwent quality control filtering. While filtering parameters were optimized for each condition, the general criteria for inclusion were as follows: 1) Mitochondrial percentage: cells with a mitochondrial percentage greater than 4% were excluded and 2) Gene count: cells with a minimum gene count of 200 and a maximum gene count ranging from 7500-10000 were retained.

These specific filtering ranges were chosen based on the distribution of each metric within the dataset to effectively remove low-quality cells, cellular debris, and potential doublets.

### Dataset normalization and integration

Gene expression data were normalized to correct for batch effects using the SCTransform method. To further refine the data, variables related to the mitochondrial percentage were regressed out. The Harmony algorithm was used for dataset integration and dimensionality reduction using the *RunHarmony* function.

### Clustering

Cells were clustered based on similar gene expression patterns using the *FindNeighbors* and *FindClusters* functions with a resolution of 0.3. To identify differentially expressed genes (DEGs) in each cluster, the *PrepSCTFindMarkers* and *FindAllMarkers* functions were used, employing the Wilcoxon Rank-Sum test. These DEGs were then used to annotate and designate the different cell types within the SC-islets. The *FeaturePlot* function was utilized to visualize the expression of specific genes across the various clusters.

### Pseudo-bulk analysis of control, IFN***α***, poly(I:C), and CVB3 conditions

The *FindMarkers* function with no logfc.threshold was used to conduct pairwise comparisons between control and treatment conditions within each time point. We then filtered the DEGs based on a significant adjusted p-value (<0.05). We used the EnhancedVolcano package to plot the DEGs.

### Analysis of single-cell populations

For the bar plots, we performed pairwise comparisons using *FindMarkers* with no logfc.threshold and filtered DEGs with a significant adjusted p-value (<0.05). We combined the time points for each treatment and used the *subset* function to isolate SC-β, -α, and -δ cells. Pairwise comparisons were performed using the *FindMarkers* function with no logfc.threshold and filtered DEGs with a significant adjusted p-value (<0.05). We used the *ggvenn* function in the ggplot2 package to create Venn diagrams. Heatmaps were generated using the pheatmap package. These DEGs were also inputted into Enrichr to perform Gene Ontology (GO) analyses.^100–102^

The PROGENy package was used to compare signaling pathway activity across different conditions and time points.^77^ First, the *progeny* function was applied to the Seurat object to calculate pathway activity scores for each cell. These scores were then scaled and centered using the *ScaleData* function. The scaled scores were transformed into a data frame and merged with cell metadata, including unique molecular identified IDs and experimental conditions. A summary table was then generated using the *summarise* function from the plyr package. This table contained the pathway names, conditions, and the mean and standard deviation of the PROGENy activity scores. Finally, a heatmap was created using the *pheatmap* function to visualize the mean PROGENy activity scores for each pathway. *pheatmap* was also used to plot the various heatmaps exhibiting various transcriptional signatures.

### Real-time quantitative PCR

Cells were harvested, washed with 1X PBS, and total RNA was extracted using the RNeasy Mini Kit (Qiagen, #74106) with on-column DNase treatment (Qiagen, #79254). Reverse transcription was performed on 100-200 ng of total RNA using the High-Capacity cDNA Reverse Transcriptase Kit (Applied Biosystems, #4368813) on a T100 Thermocycler (Bio-Rad). Real-time quantitative PCR (rt-qPCR) was conducted with PowerUp SYBR Green Master Mix (Life Technologies, #A25742) on a QuantStudio 6 Pro Thermocycler (Applied Biosystems). The relative gene expression was calculated using the ΔΔCt method. Normalization was achieved by using the average cycle threshold (Ct) of the housekeeping genes *TBP* and *GUSB*, followed by the average Ct of the corresponding genes measured from the MDA5^946T^ untreated SC-islets.

### Viability assays

Cell viability and apoptosis were assessed using a multiplexed approach combining the CellTiter-Fluor Cell Viability Assay (Promega, G6080) and the Caspase-Glo 3/7 Assay System (Promega, G8090). Briefly, SC-islets were dissociated into a single-cell suspension and plated into a 96-well, white with clear bottom plate (Corning, #3903) at a density of 4.0 x 10^4^ cells/well. After 24 h, cells were treated with either endotoxin-free water (control), 50 ng/mL IFNα, 1000 ng/mL IFNα + 500 ng/mL TNFα (R&D Systems, 210-TA), 500 ng/mL poly(I:C), or CVB3-Woodruff (titer = 1.98 x 10^12^ pfu/mL) at either a MOI of 1 or 10. 48 h post-infection, cells were washed with 1X PBS, and 80 μL of fresh culture media was added to each well. For the viability assay, 20 μL of the CellTiter-Fluor reagent was added to each well, and the plate was gently swirled. The cells were then incubated at 37°C for 30 min before measuring fluorescence with a Synergy H1 microplate reader (BioTek). Subsequently, the Caspase-Glo 3/7 assay was performed. 100 μL of the Caspase-Glo 3/7 reagent was added to each well, and the plate was incubated at room temperature in the dark for 30 min. Luminescence was then measured using the same Synergy h1 microplate reader.

### Plaque assays

Viral titers were determined using a standard plaque assay, as previously described by Burg et al. (2018).^69^

### Mitochondrial respiration assay

Mitochondrial respiration was measured using an Agilent Seahorse XFe96 extracellular flux analyzer. Briefly, SC-islets were dissociated into a single-cell suspension and plated into a Collagen IV-coated (Corning, CB40245) 96-well, Agilent Seahorse XFe96/XF Pro microplate (#103794-100) at a density of 1.0 x 10^5^ cells/well. After 24 h, cells were treated with either endotoxin-free water (control), 50 ng/mL IFNα, 500 ng/mL poly(I:C), or CVB3-Woodruff (titer = 1.98 x 10^12^ pfu/mL) at a MOI of 1. 24 h prior to the run, Seahorse XF sensor cartridges (Agilent) were hydrated with Seahorse XF Calibrant solution (Agilent, #100840-000). 48 h post-infection, the cells were washed with 1X PBS and incubated in Seahorse XF DMEM Medium (Agilent, #103575-100) supplemented with 20mM glucose, 2mM glutamine, and 1mM pyruvate at pH 7.4. The plate was then equilibrated at 37°C without CO_2_ for at least one hour. Afterward, the plate containing the SC-islets and the calibrated sensor cartridge were placed into the Seahorse XFe6 analyzer to undergo a mitochondrial stress test. Following six baseline measurements, the islets were sequentially treated with the following compounds (final concentrations in well):

1. 1.5 μM Oligomycin (Sigma, #O4876) to inhibit ATP synthase.
2. 0.5 μM Carbonyl cyanide 4-(trifluoromethoxy) phenylhydrazone (FCCP) to uncouple mitochondrial respiration.
3. 0.5 μM Rotenone (Sigma, #R8875) combined with 0.5 μM Antimycin A (Sigma, #A8674) to inhibit Complex I and III, respectively, and shut down mitochondrial respiration.

The resulting data was normalized to total DNA content per well quantified following cell lysis using the Quant-iT Picogreen dsDNA Assay Kit according to the manufacturer’s instructions.

### Statistical analysis

Statistical analyses were calculated using GraphPad Prism and R. Brown-Forsythe and Welch ANOVA test followed by Dunnett’s multiple comparisons test was used for western blot data. Two-way ANOVA followed by Sidak’s multiple comparisons test as well as an unpaired Student t-test were used for flow cytometry data. Multiple unpaired t-tests and two-stage linear step-up procedure of Benjamini, Krieger, and Yekutieli, as well as RM two-way ANOVA followed by Dunnett’s multiple comparisons test were performed on sGSIS data. Welch’s unpaired t test and ordinary one-way ANOVA with Dunnett’s multiple comparisons test were used for INS content data. Wilcoxon rank-sum test in conjunction with Bonferroni correction were used to obtain statistically significant DEGs with an adjusted *p* value < 0.05. We used either ordinary one-way ANOVA or Brown-Forsythe and Welch ANOVA tests, followed by Sidak’s and Dunnett’s multiple comparisons test, respectively, for real-time qPCR (rt-qPCR). The rt-qPCR for *VP1* used an unpaired t-test. Ordinary two-way ANOVA followed by Sidak’s multiple comparisons test was used for caspase-3/7 assay data. An unpaired t-test with Welch’s correction was used for plaque assay data.

The sample sizes for flow cytometry and sGSIS data contained four differentiation batches for each variant. Three differentiation batches were used for INS content, rt-qPCR, and caspase 3/7 data. The sample sizes for scRNA-seq and OCR measurements contained two separate SC-islet differentiation batches per variant. To measure intracellular viral levels, three differentiation batches were used for MDA5^946T^ and MDA5^627^*. We used GraphPad Prism to identify any outliers using the Grubbs’ method in our plaque assays. All statistical details can be found in the figure legends. *p* values are marked as follows: non-significant (ns) *p* > 0.05, ^∗^*p* < 0.05, ^∗∗^*p* < 0.01, ^∗∗∗^*p* < 0.001, and ^∗∗∗∗^*p* < 0.0001.

## Author contributions

D.A.V-P. and J.R.M. designed all experiments and wrote the manuscript. D.A.V-P performed all sequencing and computational experiments. D.A.V-P., C.B., K.B., N.M., S.E.G., and K.E.H. performed all *in vitro* experiments. C.E.M. provided the CRISPR-Cas9-edited CH1-064 hPSCs. H.M.T. and C.E.M. provided key guidance, insights, and reagents. All authors revised, reviewed, and approved of the manuscript.

## Acknowledgements

This work was primarily funded by the National Institutes of Health (NIH) (R01DK138469) to J.R.M. and H.M.T., with additional support from NIH (R01DK114233, R01DK127497, and R01DK126456), Breakthrough T1D (3-SRA-2023-1295-S-B), the Edward J. Mallinckrodt Foundation, and Washington University School of Medicine Department of Medicine startup funds to J.R.M. D.A.V.-P. was supported by the National Science Foundation’s (NSF) Graduate Research Fellowship Program (DGE 2139839). Further support came from the Washington University Diabetes Research Center (P30DK020579). We thank the Genome Technology Access Center at the McDonnell Genome Institute at Washington University School of Medicine for genomic analysis support. The center is partially supported by NCI Cancer Center Support Grant P30CA91842 to the Siteman Cancer Center from the National Center for Research Resources (NCRR), a component of the NIH, and NIH Roadmap for Medical Research. This publication is solely the authors’ responsibility and does not represent the official views of NIH, NSF, or any other funder. We would also like to thank Erika Brown (Washington University) for helpful feedback on the manuscript.

