## Extended data figures for "Protective *IFIH1* variant reduces immune-mediated islet stress and dysfunction in a type 1 diabetes genetic background"

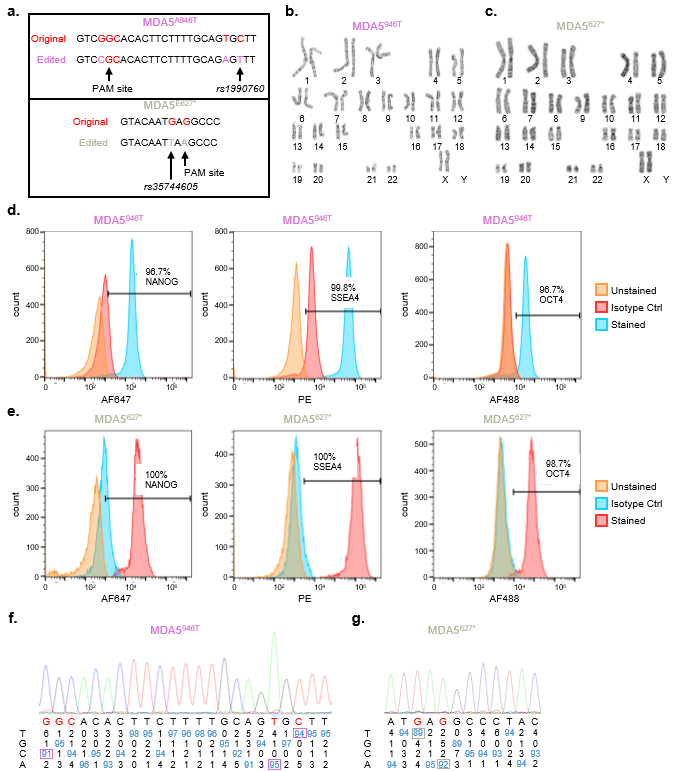

Extended Data Figure 1. Characterization of CRISPR-Cas9-edited hPSCs. (A), Schematic depicting CRISPR-Cas9 editing of *IFIH1* at different loci to generate hPSCs harboring either the MDA5^946T^ or MDA5^627*^ variants. (B-C), Normal 46XY karyotype of CRISPR-Cas9-edited (B) MDA5^946T^ and (C) MDA5^627*^ hPSCs. (D), Quantification of pluripotent protein marker expression derived from flow cytometry measurements using MDA5^946T^ hPSCs. (E), Quantification of pluripotent protein marker expression derived from flow cytometry measurements using MDA5^627*^ hPSCs. (F-G), Sanger sequencing of (F) MDA5^946T^ and (G) MDA5^627*^ hPSCs to confirm proper CRISPR-Cas9 editing.

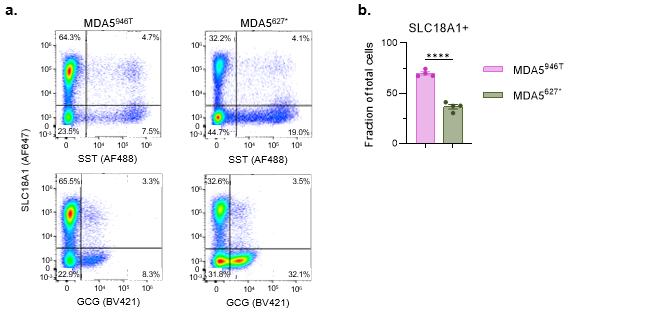

Extended Data Figure 2. *IFIH1* variant-derived SC-islets exhibit differing amounts of off-target populations. (A), Representative flow cytometry dot plots of dispersed SC-islets for each MDA5 variant for SLC18A1, SST, and GCG. (B), Quantified fraction of cells expressing SLC18A1 for SC-islets derived from each MDA5 variant (n = 4). Error bars represent s.e.m. ** = Unpaired t test.

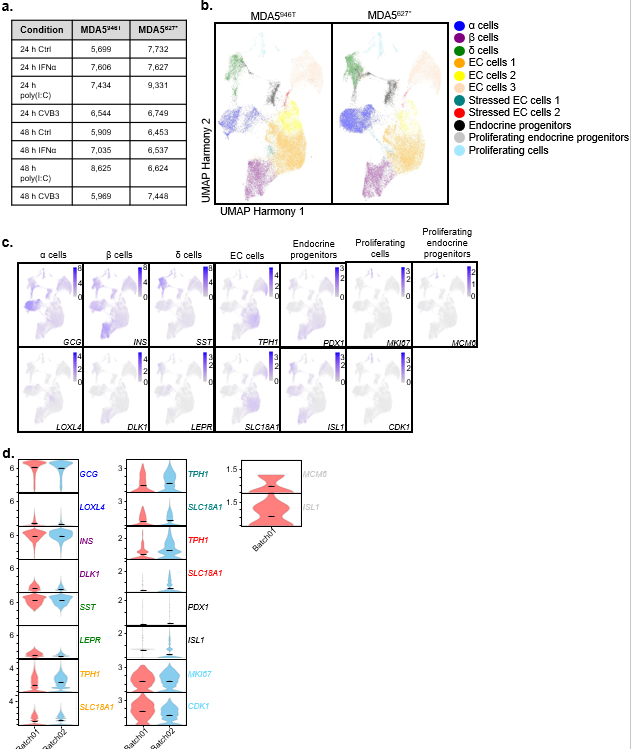

Extended Data Figure 3. scRNA-seq similarly captures SC-islet-associated cell types across MDA5 variants and differentiation batches. (A), Cell numbers obtained from each condition and time point for scRNA-seq following quality check and filtering. (B), Harmony UMAPs depicting cluster distribution across MDA5 variants. (C), Feature plots showing cell marker expression within clusters. (D), Violin plots exhibiting cell marker expression across differentiation batches.

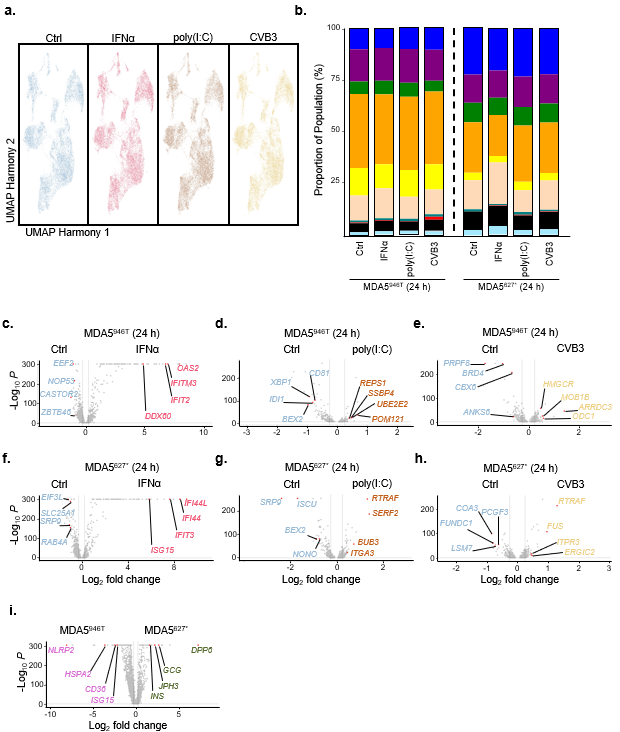

Extended Data Figure 4. Single-cell RNA sequencing reveals major transcriptional differences between control and treated SC-islets. (A), Harmony UMAPs showing cluster distribution across stress conditions. (B), Stacked bar plot depicting the proportion of cells found in SC-islets derived from each MDA5 variant and treated with various stressors for 24 h. (C-H), Volcano plots showing transcriptional differences when comparing (C) MDA5^946T^ Ctrl vs IFNα 24 h (total variables = 2066), (D) MDA5^946T^ Ctrl vs poly(I:C) 24 h (total variables = 382), (E) MDA5^946T^ Ctrl vs CVB3 24 h (total variables = 1197), (F) MDA5^627*^ Ctrl vs IFNα 24 h (total variables = 1880), (G) MDA5^627*^ Ctrl vs poly(I:C) 24 h (total variables = 940), (H) MDA5^627*^ Ctrl vs CVB3 24 h (total variables = 872). (I) Volcano plot showing transcriptional differences between whole SC-islets from *IFIH1* variants at baseline (total variables = 3983).

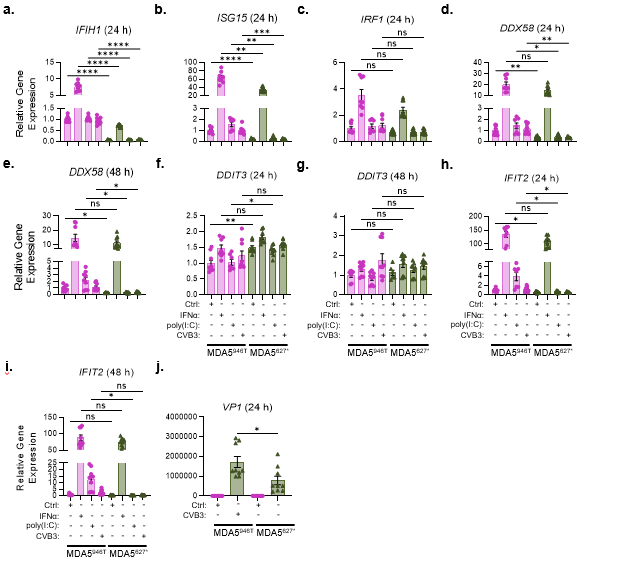

Extended Data Figure 5. rt-qPCR reveals differences in stress response between MDA5 variants. (A-I), rt-qPCR of type 1 IFN-associated genes. (J), rt-qPCR of viral genome expression in CVB3-infected SC-islets. Error bars represent s.e.m. ** = Brown-Forsythe and Welch ANOVA test followed by Dunnett’s multiple comparisons test (A-F, H-I), ordinary one-way ANOVA followed by Sidak’s multiple comparisons test (G), unpaired t test (J).

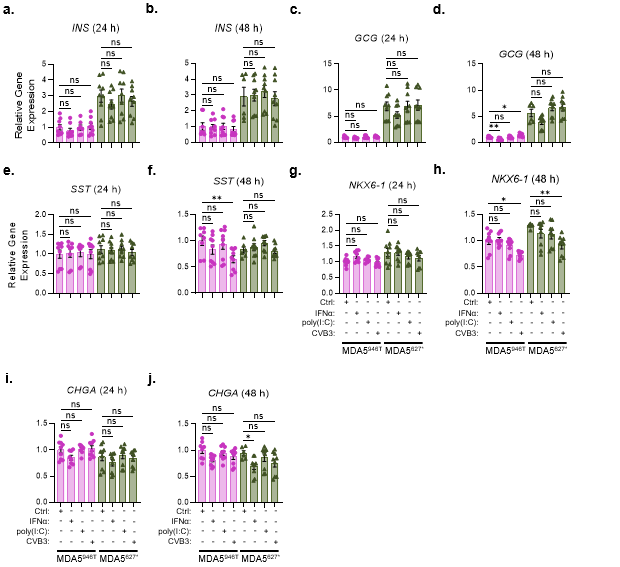

Extended Data Figure 6. rt-qPCR reveals no differences in islet identity response to stress between MDA5 variants. (A-J), rt-qPCR of islet identity genes. Error bars represent s.e.m. ** = Brown-Forsythe and Welch ANOVA test followed by Dunnett’s multiple comparisons test (A-D, G-H), ordinary one-way ANOVA followed by Sidak’s multiple comparisons test (E-F, I-J).
